# Timing of neurogenesis through sequential accumulation of miR-9 due to additive expression of multiple alleles

**DOI:** 10.1101/2020.08.03.233890

**Authors:** X. Soto, J. Burton, C. Manning, T. Minchington, R. Lea, J. Lee, J. Kursawe, M. Rattray, N. Papalopulu

## Abstract

MicroRNAs (miRs) have important quantitative roles in tuning dynamical gene expression. Hes/Her transcription factor dynamics are sensitive to the increasing amount of miR-9 in the cell, transitioning from noisy high-level expression to oscillatory expression and then to downregulation. However, the mechanism by which miR-9 is quantitatively controlled is not known. In vertebrates, several distinct genomic loci produce the same mature miR-9, but the functional significance of multiple primary transcripts remains unknown. Here, we show that the amount of mature miR-9 increases during zebrafish neurogenesis in a sharp stepwise manner. We characterize the spatiotemporal profile of 7 distinct pri-mir-9s and show that they are sequentially expressed during hindbrain neurogenesis. Quantitative analysis of expression at the single-cell level, shows that expression of late-onset pri-mir-9-1 is added on, rather than replacing the expression of early onset pri-mir-9-4 and 9-5. Mutating the late-onset pri-mir-9-1 with CRISPR/Cas9 prevents the developmental increase of mature miR-9 and reduces late neuronal differentiation. Finally, we use mathematical modelling to explore possible benefits of a stepwise increase of miR-9 over a linear increase. We find that an adaptive network containing Her6 can be insensitive to a linear increase in miR-9 and show that such adaptation can be overcome by step-wise increases of miR-9. In conclusion, our work suggests that a sharp stepwise increase of mature miR-9 is contributed by sequential temporal activation of distinct loci. This may be a strategy to overcome adaptation and facilitate a transition to a new state of Her6 dynamics or level.

## Introduction

MicroRNAs are a class of small (∼22 nucleotides) regulatory non-coding RNAs, which regulate gene expression at the post-transcriptional level. These small RNAs are processed from large microRNA primary transcripts (pri-mir) into 70∼90nt precursors (pre-mir) before further splicing into ∼22nt mature microRNA (miR). miR-9 is a highly conserved microRNA that is expressed predominantly in the central nervous system (CNS) of vertebrates and plays a crucial role during CNS development. Specifically, previous work in Xenopus, zebrafish and mice has shown that miR-9 is essential for cell fate transitions during neurogenesis (Shibata et al., 2011) (Coolen et al., 2013) (Bonev et al., 2011; Bonev et al., 2012). miR-9 post-transcriptionally targets many transcription factors that are involved in neural development such as FoxG1 (Shibata et al., 2008), Tlx (Zhao et al., 2009) and members of the Hes/Her helix-loop-helix family of transcription factors, including Hes1 in mouse and Xenopus (Bonev et al., 2011; Bonev et al., 2012) and Her6/Her9 in zebrafish (Coolen et al., 2012) (Galant et al., 2016) (Soto et al., 2020) (Leucht et al., 2008).

The Hes/Her family of proteins are expressed dynamically in an oscillatory manner at the ultradian timescale (Hirata et al., 2002) (Shimojo et al., 2008). Hes/Her oscillations are achieved by a negative feedback loop, whereby Hes/Her proteins inhibit their own transcription coupled with a rapid turnover of protein and mRNA. Instability of both protein and mRNA allows for levels of the protein to fall, de-repression to occur and expression to resume, generating a cyclic pattern (Hirata et al., 2002) (Novak and Tyson, 2008). Indeed, both mRNA and protein of Hes family genes are unstable, for example, in mice, the half-life of *Hes1* mRNA is ∼24 minutes; the Hes1 protein half-life is in the order of 22 minutes (Hirata et al., 2002) and the Her6 (Hes1 Zebrafish orthologue) protein half-life is around 10 minutes (Soto et al., 2020).

Instability of mRNA, as well as translation of protein, are partly controlled by microRNAs. Indeed, our previous work revealed that miR-9 regulation is important for controlling *Hes1* mRNA stability and allowing the oscillatory expression of Hes1 to emerge. We have recently shown that in zebrafish, the dynamics of Her6 protein expression switch from noisy to oscillatory and then to downregulation and that these changes coincide temporally with the onset of miR-9 expression in the hindbrain (Soto et al., 2020). When the influence of miR-9 on *her6* is removed experimentally, Her6 expression does not evolve away from the “noisy” regime and is not downregulated with a consequent reduction in progenitor differentiation. We have interpreted this to mean that the miR-9 input is necessary to constrain gene expression noise, enabling oscillations to occur and to be decoded by downstream genes which in turn participate in downregulating Her6 as cells differentiate (Soto et al., 2020).

However, not only the presence of miR-9 but also the amount of miR-9 present is important, as too much or too little miR-9 can lead to dampening of Hes1 oscillations (Bonev et al., 2012) (Goodfellow et al., 2014). Indeed, mathematical modelling showed that increasing miR-9 over time drives the Hes1 expression into different states (oscillatory or stable high/low) and that the amount of miR-9 present in the cell determines the length of time by which Hes1 oscillates, effectively timing the transition to differentiation (Phillips et al., 2016) (Goodfellow et al., 2014). Together these findings support that Hes/Her dynamics and downregulation are sensitive to the amount of mature miR-9 present in the cell; however, the mechanism by which miR-9 level is controlled is not known.

This question is complicated by the observation that vertebrates (and some invertebrates) possess multiple copies of the miR-9 gene at distinct loci which are all capable of producing the same mature microRNA. For example, both human and mouse contain 3 copies of miR-9 (Rodriguez-Otero et al., 2011; Shibata et al., 2011) and frogs have 4 (Walker and Harland, 2008). Due to an additional round of whole-genome duplications in teleost fish (Amores et al., 1998) (Jaillon et al., 2004), zebrafish have 7 paralogues of miR-9 (pri-mir-9-1 to pri-mir-9-7) (Chen et al., 2005).

One possibility is that different genomic loci contribute to miR-9 regulation in a qualitative way, with differential temporal and spatial specificity of mature miR-9 expression. Indeed, there is some limited evidence that these discrete copies of miR-9 are expressed differentially during development both temporally and spatially (Nepal et al., 2016; Tambalo et al., 2020). Another, and yet unexplored, possibility is that transcription from different loci may serve to control miR-9 quantitatively, that is to increase the amount of miR-9 in the cell and perhaps do so in a temporally controlled manner, thus, contributing to the change of miR-9 levels that is necessary to drive a change in the dynamics of Hes/Her targets.

Here, we undertake a systematic study of pri-mir-9 expression in zebrafish that aims to address the likelihood of these distinct scenarios, with special attention to the possibility of a quantitative control mechanism. We show by *in situ* hybridization that the expression of miR-9 spreads from the forebrain to the hindbrain and increases quantitatively in the hindbrain between 24 and 48hpf. A detailed time course of the expression of all 7 pri-mir-9 paralogues shows that they are all transcriptionally active, but exhibit subtle, yet distinct, temporal and spatial profiles. Focusing on a set of early and late expressed pri-mir-9s in the hindbrain (pri-mir-9-1, pri-mir-9-4 and pri-mir-9-5) by quantitative single molecule fluorescent in situ hybridisation at single cell level, we find that, surprisingly, in many cells, early and late pri-mir-9s were concurrently transcriptionally active such that the expression from late activated pri-mir-9s is added on to the early ones. This is functionally significant as the specific mutation of the late pri-mir-9-1 selectively reduces neurons that normally differentiate late. Taken together, we find evidence for a qualitative mechanism in the deployment of pri-mir-9s, as well as a previously un-appreciated temporally controlled quantitative component. While both quantitative and qualitative mechanisms may contribute to the decoding function of mature miR-9s, our mathematical modelling suggests the sharp quantitative increase afforded by the deployment of additional transcriptional units, may facilitate the downregulation of Her6 at late time points.

## RESULTS

### Pri-mir-9s are expressed with differed temporal onset

miRs are derived from a duplex precursor and the -5p strand (“guide”) is preferentially incorporated into an RNA-induced silencing complex (RISC) to exert its regulatory functions, while the complementary -3p strand (“passenger”) is thought to be rapidly degraded. Indeed, for the mature miR-9 the miR-9-5p is designated as the “guide” strand and its annotation is derived from the mature miR sequence being embedded in the 5′ stem of the miR-9 precursor.

To investigate the expression of the mature miR-9 (−5p strand) in zebrafish embryos, we first performed a whole-mount in situ hybridization (WM-ISH) for the mature miR-9 using an LNA (locked nucleic acid) probe. Mature miR-9 was detected only in the forebrain at 24hpf (**Fig. 1A**) while at 30hpf miR-9 is weakly observed in the midbrain and rhombomere 1 (r1) of the hindbrain (hb), maintaining high expression in the forebrain (**Fig. 1A**, 30hpf). As development progressed, miR-9 expression increased in the hindbrain with steady high levels in the forebrain (**Fig. 1A**, 34-36hpf-blue arrow). But later in development levels in the hindbrain was further increased while in the forebrain were decreased (**Fig. 1A**, 48hpf, blue arrow: hindbrain, green arrow: forebrain). These results show a temporally controlled increase of miR-9 expression along the brain/hindbrain axis as previously described in (Soto et al., 2020).

**Figure 1.**
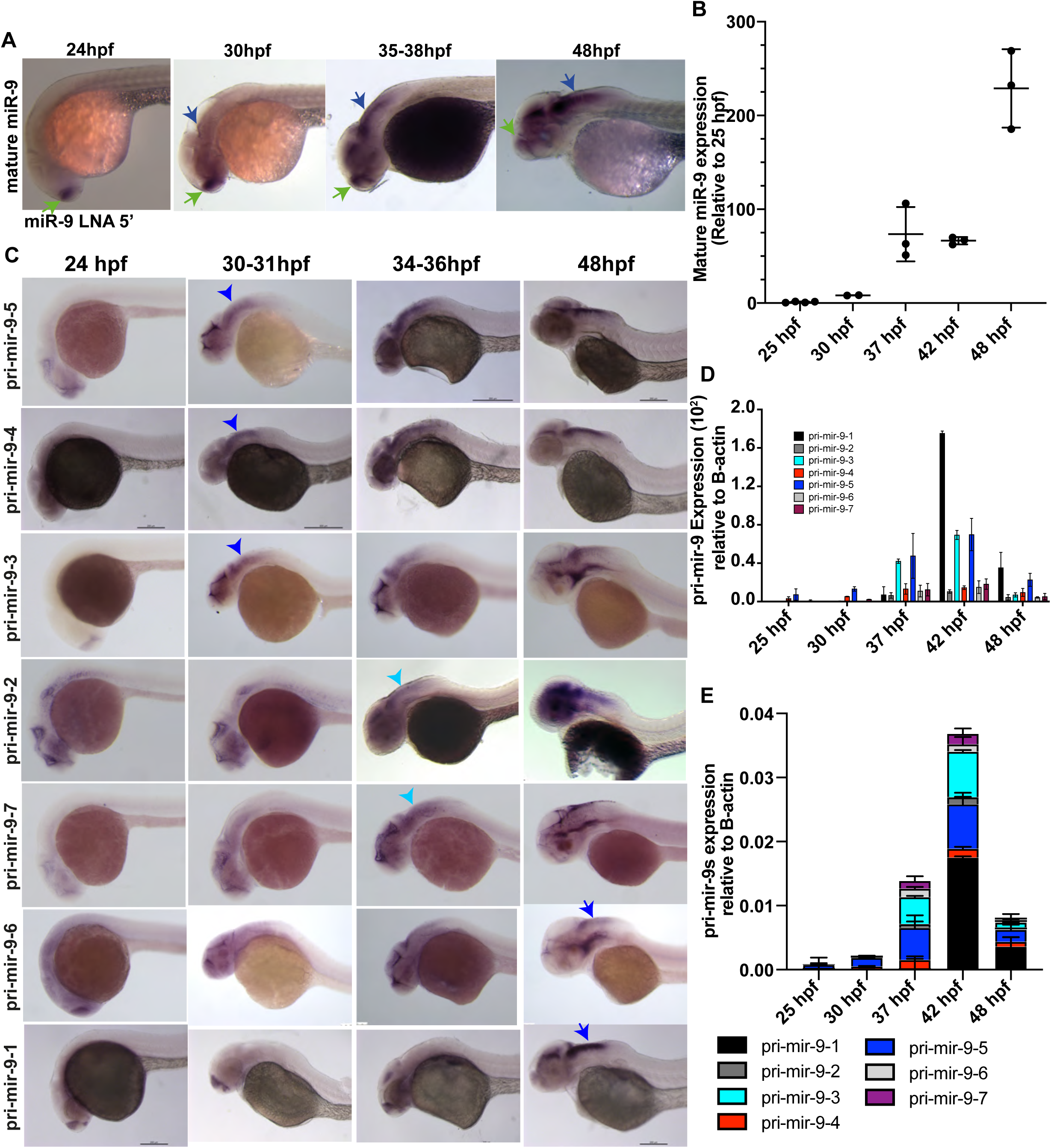
Pri-mir-9 paralogues are expressed with different temporal onset. **(A)** Representative example of chromogenic whole mount in situ hybridization (WM-ISH) of miR-9 using miR-9 LNA 5’-Dig observed at different stages during development; Similar results can be observed in (Soto et al., 2020). Longitudinal view, anterior to the left. Green arrow: forebrain (fb) expression. Blue arrow: hindbrain (hb) expression. **(B)** Taqman RT-qPCR of mature miR-9 from dissected hindbrain at different stages of development, relative to 25hpf. **(C)** Chromogenic WM-ISH (Cro-WMISH) of different pri-mir-9s using specific probes for each paralogue observed at different stages during development; longitudinal view, anterior to the left. Blue arrowhead: expression in hb at 30-31hpf, light blue arrowhead: expression in hb at 34-36hpf, blue arrow: expression in hb at 48hpf. **(D**,**E)** SYBR green RT-qPCR relative quantification of the seven pri-mir-9s from dissected hindbrain at different stages of development, quantification was normalised using β-actin. **(B, D, E)** N=3, each N contain a pool of 10 hindbrains.

To characterize the increase of expression in the hindbrain in a quantitative manner, we used RT-qPCR on dissected hindbrains from stages 25hpf to 48hpf. This analysis confirmed that there is upregulation in the time frame analysed. An initial low level of expression at 30hpf was followed by a sharp upregulation at 37hpf, which was maintained through to 42hpf, undergoing a second sharp increase at 48hpf **(Fig. 1B)**.

In zebrafish, the mature miR-9 can be produced from seven paralogues of miR-9. The miR-9 paralogues occupy 7 unique loci across the genome (GRCz11; Genome Reference Consortium Zebrafish Build 11) (Yates et al., 2020). With the exception of miR-9-3 which is located upstream of a lincRNA, all miR-9 genes are intragenic, overlapping annotations of lincRNAs or proteins (Yates et al., 2020) (**Fig. S1A**,**B**). Our *in silico* analysis of previously published RNA-seq data shows differential temporal expression of six of the seven miR-9 paralogues hosts (White et al., 2017). It is also clear that upregulation of miR-9 host genes (and hence miR-9) coincides with a gradual decline in the expression of Her/Hes family genes expression, consistent with the idea that *hes/her* genes are major targets of miR-9 (**Fig. S1C**) (Bonev et al., 2011).

Previous work has revealed that the 7 miR-9 zebrafish paralogues are expressed in forebrain at early stages of neurogenesis moreover toward the end of embryonic neuronal differentiation they are also expressed in hindbrain (Nepal et al., 2016). However, little is known about the period spanning the peak of neurogenesis, when miR-9 controls downstream targets such as the ultradian oscillator Her6. To characterize the expression in greater spatiotemporal detail, particularly over regions of the hindbrain area where Hes/Her target genes are expressed, we investigated the expression of all 7 primary transcripts over a time period spanning the peak of neuro-genesis which occurs at 33hpf (Lyons et al., 2003) using specific probes for each pri-mir-9 (**Fig. S2; Methods**, Molecular cloning).

We observed that all pri-mir-9s were first expressed in the forebrain (24hpf), in a regional specific manner, which was not further characterised here. At 48hpf they are all also expressed in hindbrain (**Fig. 1C**, 24 and 48hpf and **Fig. S3A-C**) consistent with what has been described before (Nepal et al., 2016). Differential expression was evident in the intermediate stages. Specifically, pri-mir-9-3, 9-4 and 9-5 were expressed ahead of the others in the hindbrain (**Fig. 1C**; 30hpf; blue arrowhead). At the peak of hindbrain neurogenesis (34-36hpf), pri-mir-9-2 and 9-7 were upregulated, joining most of the pri-mir-9s that were highly expressed at this stage (**Fig. 1C**, 34-36hpf). Pri-mir-9-1 and 9-6 were temporally delayed showing hindbrain expression at 48hpf, at which point all pri-mir-9 were fully expressed. Quantifying the expression with RT-qPCR confirmed that pri-mir-9-4 and 9-5 were expressed early and that expression of pri-mir-9-1 commenced relatively late, at 42hpf. At 48hpf, all pri-mir-9s had lower level of expression although pri-mir-9-1 continued to be relatively high compared to the other pri-mir-9s (**Fig. 1D and E**). Overall, every pri-mir-9 was expressed in the CNS and exhibited a temporal progression.

### Expression of pri-mir-9s from distinct loci is additive and sequentially activated

To achieve a more detailed characterisation of expression, we selected three different primary transcripts based on; (1) the onset of their hindbrain temporal expression during development, earliest or latest, and (2) a phylogenetic analysis of sequence based on vertebrates evolutionary relationship performed by (Alwin Prem Anand et al., 2018), to select representatives that are widely distributed in the phylogenetic tree. Thus, pri-mir-9-5 was selected as the earliest to be expressed in the hindbrain and belonging to clade I/subgroup I, pri-mir-9-4 as the earliest and belonging to clade II and pri-mir-9-1 as the latest and belonging to clade I/subgroup II (Alwin Prem Anand et al., 2018) (**Fig. S3D)**.

Double WM-FISH for pri-mir-9-1/pri-mir-9-4 and pri-mir-9-1/pri-mir-9-5 performed on stage 30-32hpf embryos revealed expression of pri-mir-9-4 and pri-mir-9-5 along the anterior-posterior (A-P) hindbrain axis, while the expression of pri-mir-9-1 at this early stage was limited to the region of the anterior hindbrain corresponding to rhombomere 1 (**Fig. 2A**, red arrow). A transversal view at posterior hindbrain (rhombomere 4) reveals expression of pri-mir-9-4 and pri-mir-9-5 within the ventricular zone (VZ; **Fig. 2A-B,E**) indicating that pri-mir-9s are expressed in the region where most of the progenitors are found (Lyons et al., 2003; Tambalo et al., 2020). Pri-mir-9-1 staining shows an artefactual surface expression, as indicated with white arrow in transversal view at 30-32hpf, this is produced due to the WM-FISH detection method.

**Figure 2.**
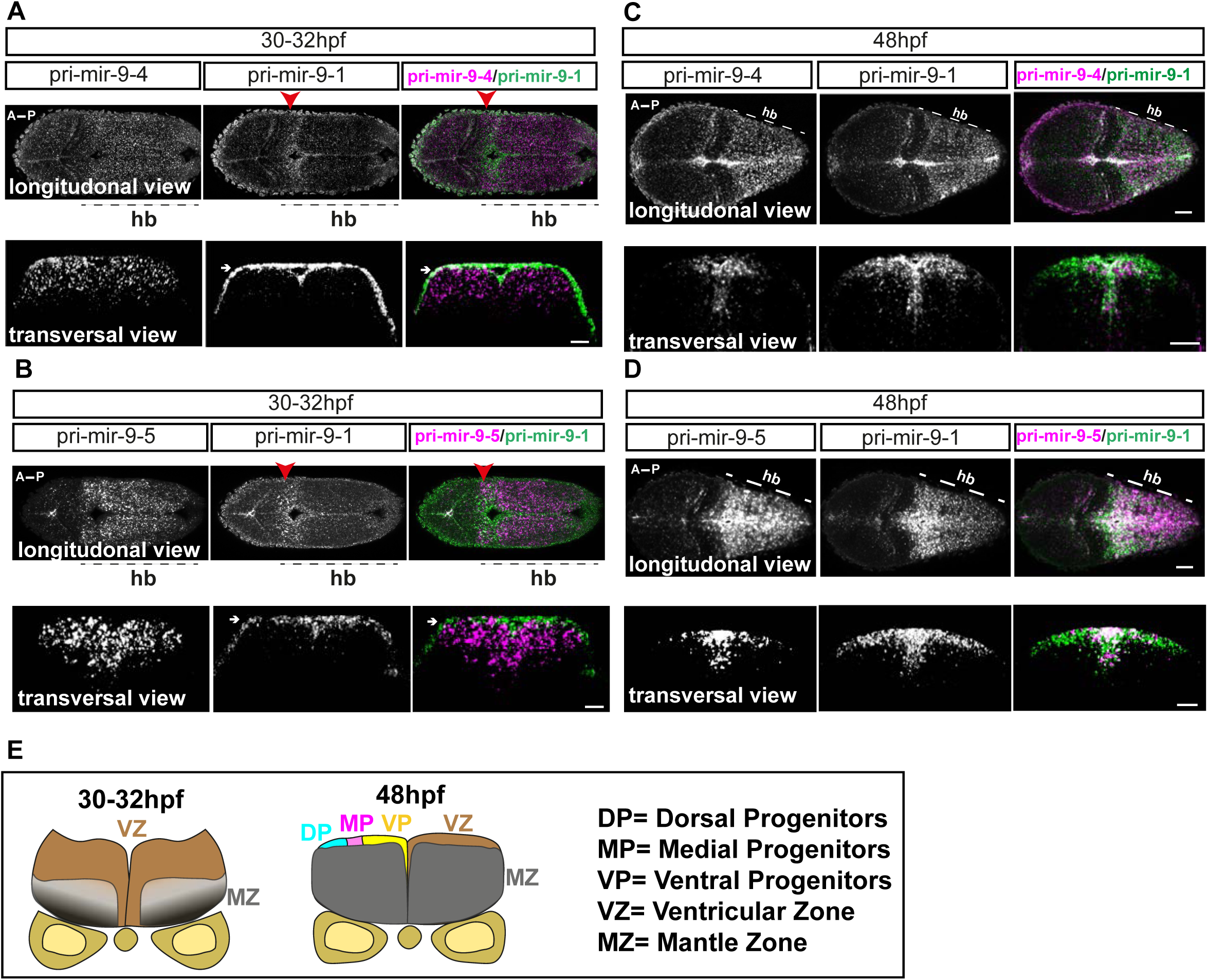
Progressive additive expression of pri-mir-9s during development. **(A-D)** Representative example of double Fluorescent WM-ISH (WM-FISH) labelling pri-mir-9-1/pri-mir-9-4 **(A,C)** and pri-mir-9-1/pri-mir-9-5 **(B,D)** in hindbrain (hb) from wild-type embryo observed at 30-32hpf **(A,B)** and at 48hpf **(C,D). Transversal view** was collected from hindbrain rhombomere 4/5 (r4/5). **Longitudinal view** was collected from embryos with anterior to the left and posterior to the right; images are maximum intensity Projection; 5 microns thickness for 48hpf embryos and 10 microns thickness for 30-32hpf embryos. Merged images indicate pri-mir-9-4 or 9-5 in magenta and pri-mir-9-1 in green. White arrows indicated artefactual signal originated from the amplification step with FITC staining in the WM-FISH Red arrows indicate rhombomere 1 (r1) of hb. Scale bar 30 μm. Annotations indicate **A**: anterior, **P:** posterior, hb: hindbrain. (**N pri-mir-9-1/9-4** = longitudinal/30-32hpf: 3; transversal/ 30-32hpf: 3; longitudinal/48hpf: 4; transversal/48hpf: 8); (**N pri-mir-9-1/9-5** = longitudinal/30-32hpf: 3; transversal/ 30-32hpf: 4; longitudinal/48hpf: 4; transversal/48hpf: 5). **(E)** Schematic representation of transverse section from zebrafish hindbrain at the level of the otic vesicle for 30-32hpf and 48hpf, respectively. VZ: ventricular zone, region where most of progenitor cells are located. MZ: mantle zone, region of ongoing neurogenesis. Within the VZ there are dorsal progenitors (DP), medial progenitors (MP) and ventral progenitors (VP).

We repeated this analysis at 48hpf to examine whether the late expression of pri-mir-9-1 is cumulative with pri-mir-9-4 and pri-mir-9-5 or spatially distinct. Double WM-FISH of pri-mir-9-1 with pri-mir-9-4 or pri-mir-9-5 revealed overlapping expression of the primary transcripts in both longitudinal and transversal views (**Fig. 2C, D**). In addition, some distinct expression was observed in transversal views in that pri-mir-9-1 was more broadly expressed toward the dorsal progenitor region (**Fig. 2C-E**) when compared to pri-mir-9-4 and pri-mir-9-5, respectively.

### Mature miR-9 accumulates in single cells by overlapping expression of distinct loci primary transcripts

For overlapping expression to contribute to the total levels of mature miR-9 in a cell, early and late pri-mir-9s would need to be expressed in the same cells. Thus, we investigated pri-mir-9 expression at the single-cell level, by triple WM-smiFISH for pri-mir-9-1, 9-4 and 9-5, to detect nascent transcription sites, and Phalloidin staining to reveal cell boundaries. At 30hpf we observed that most cells expressed only one miR-9 primary transcript, pri-mir-9-4 or pri-mir-9-5, while a small proportion expressed both and none expressed pri-mir-9-1 (**Fig. 3A, D-F)**. By contrast at 36-37hpf and 48hpf, the number of cells that expressed one pri-mir-9 decreased and correspondingly, the number that expressed 2 or 3 pri-mir-9s increased. The most striking increase was observed in the number of cells that co-express 3 pri-mir-9s at 36-37hpf, which was due to the onset of transcription of pri-mir-9-1 in the same cells that expressed pri-miR-9-4 and 9-5. This finding suggested that in many hindbrain cells the late expression of pri-mir-9-1 is added to the earlier expression of pri-mir-9-4 and 9-5.

**Figure 3.**
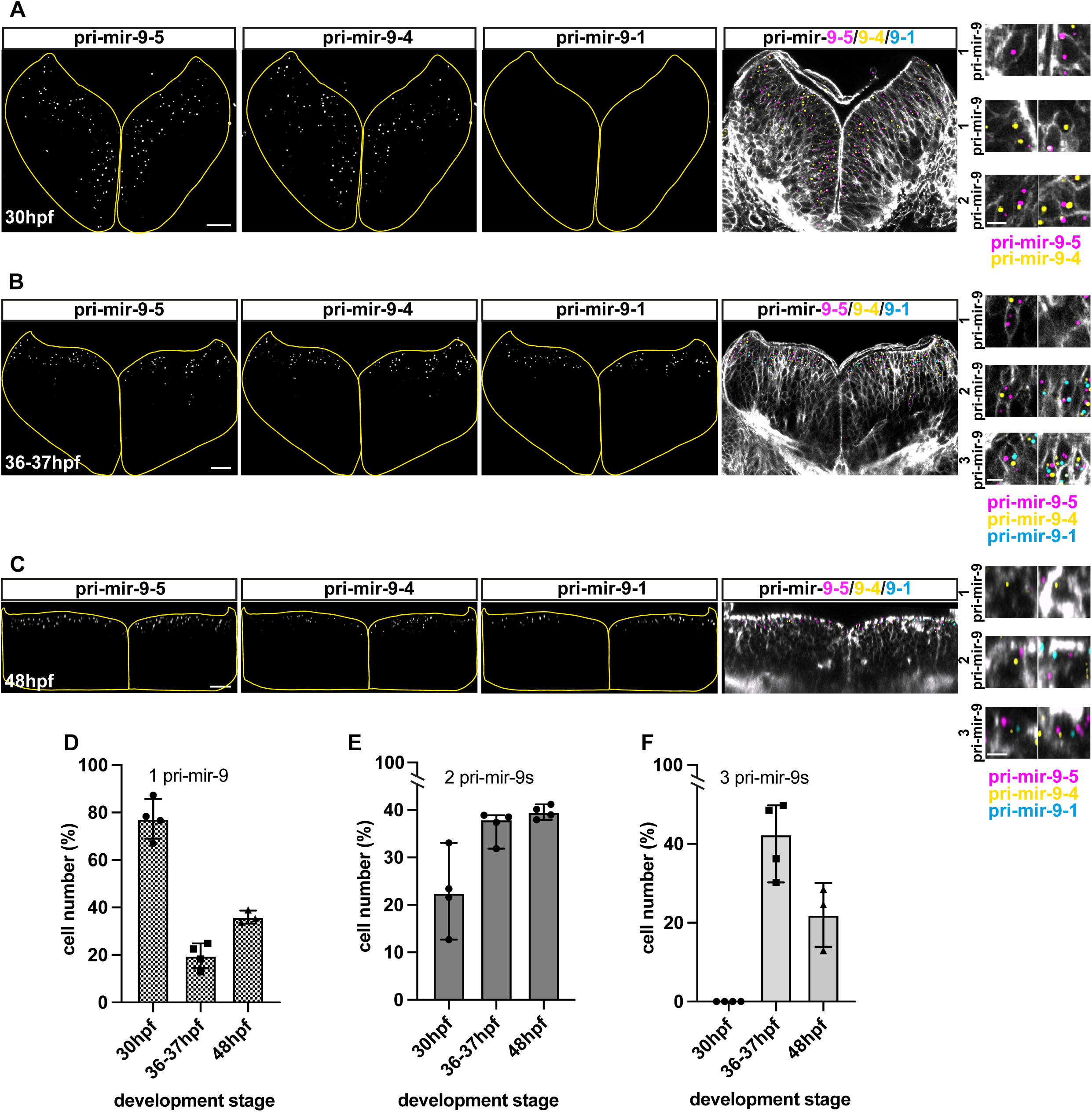
mature miR-9 expression in a cell is contributed by overlapping activation of distinct miR-9 loci. **(A-C)** Representative example of transversal view from triple whole mount smiFISH labelling active transcriptional sites for pri-mir-9-5, -9-4 and -9-1 (from left to right) combined with cell boundary staining, Phalloidin-Alexa Fluor 488, (observed in gray color in merge images) in hindbrain from wild-type embryo at 30hpf **(A)**, 36-37hpf **(B)** and 48hpf **(C)**. Merged images indicate pri-mir-9-5 in magenta, pri-mir-9-4 in yellow, pri-mir-9-1 in cyan and membrane in gray. Scale bar 20μm. **(A’-C’)** Increased magnification of representative images to show single cells expressing any single pri-miR-9 (**1 pri-mir-9)**, any two different pri-miR-9 **(2 pri-mir-9)** and the three different pri-miR-9 **(3 pri-mir-9)**. Scale bar 5μm. **(D-F)** Percentage of cells expressing any single pri-mir-9 **(D)**, any two different pri-miR-9 **(E)** and the three different pri-miR-9 **(F)** expressed relative to total number of cells positive for the precursors. N 30hpf= 4; N37-37hpf= 4; N 48hpf= 3

### Medial and Dorsal progenitor maintain concurrent expression of miR-9 primary transcripts at late neurogenesis

Based on the smiFISH data presented above we created a map that depicts transcription in single cells in transverse sections of the hindbrain over development (**Fig 4A-C**). From left to right we observe **(i)** the whole transverse section obtained from the Phalloidin staining, **(ii)** the region where cells transcribe pri-mir-9-5, **(iii)** cells with overlapping transcription for pri-mir-9-5/9-4 and **(iv)** the cells in which the three primary transcripts are transcribing (**Fig. 4A-C**). At 30hpf, pri-mir-9-4 and 9-5 are co-expressed in many cells of the ventricular zone (VZ) (**Fig. 4A**). At 36-37hpf, pri-mir-9-1 is transcriptionally activated in most, but not all, neural progenitors that already express pri-mir-9-4 and 9-5 (**Fig. 4B**) and at 48hpf the pattern of triple pri-mir-9 co-expression is maintained as similar as in 36-37hpf (**Fig. 4C**). This result supports the concurrent expression of pri-mir-9s at late stages but also shows their expression in the neural progenitor area, which thins during development as cell differentiate (**Fig. 4D**). All 3 paralogues are switched off in differentiating cells located in the Marginal Zone suggesting that they are involved in the decision to differentiate rather than maintaining the differentiated state (**Fig. 4A-C**).

**Figure 4.**
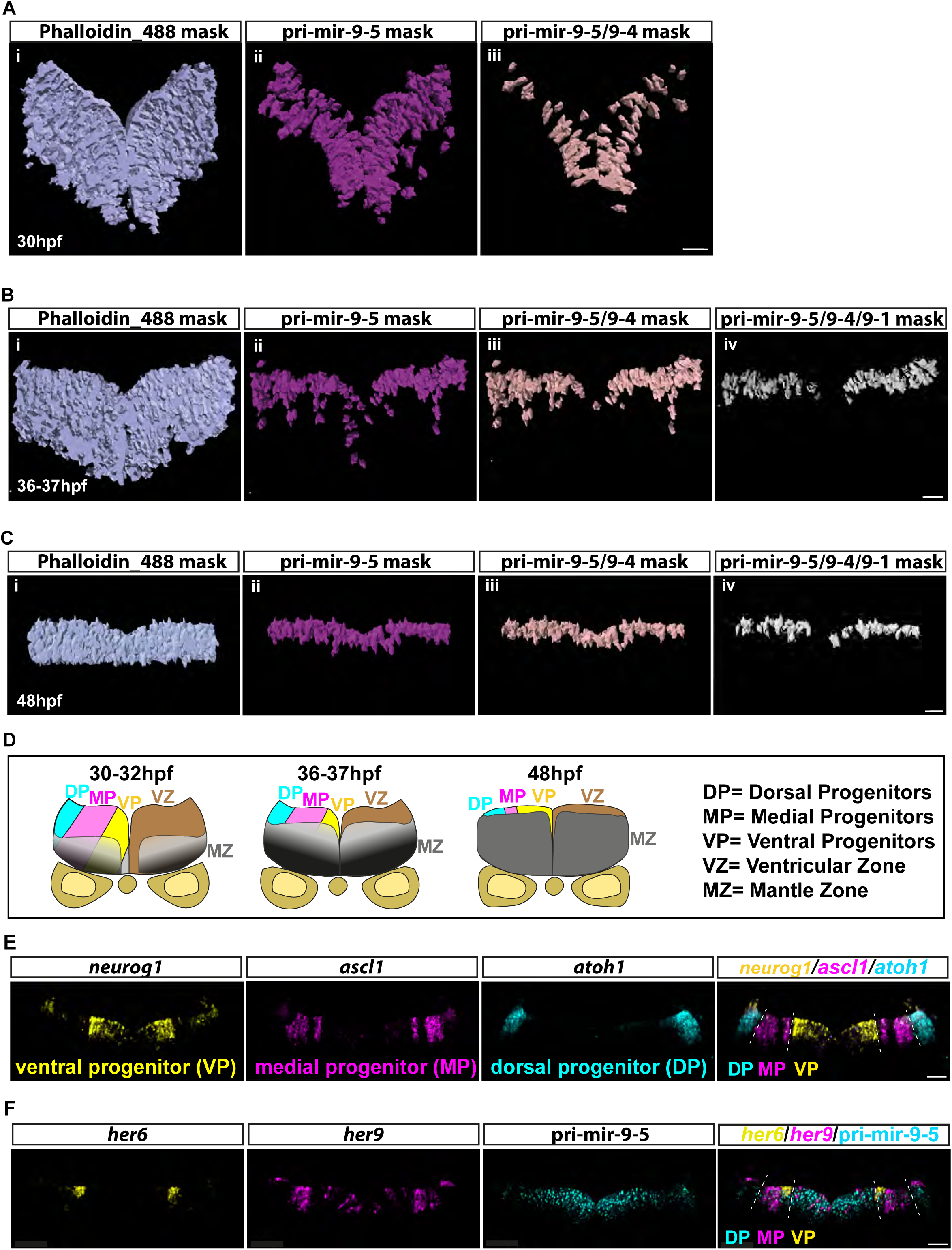
Concurrent expression of miR-9 precursors in dorsal and medial progenitors. **(A-C)** Mask representing segmented cells obtained from confocal images in **Figure 3**. Using Imaris software, the cell segmentation was performed based on the membrane marker, Phalloidin-AF488, and the spot tool allowed us counting active transcriptional sites for pri-mir-9-5, -9-4 and -9-1, respectively. From left to right we visualize the mask showing all segmented cells present in the transversal view of the hindbrain **(i, light blue)**, segmented cells that express pri-mir-9-5 **(ii, magenta)**, both pri-mir-9-5 and -9-4 **(iii, light pink)** and all three pri-mir-9-5, -9-4 and - 9-1 **(iv, gray)**. The study was performed at 30hpf **(A)**, 36-37hpf **(B)** and 48hpf **(C)**. Scale bar 20μm. **(D)** Schematic representation of transverse section from zebrafish hindbrain at the level of the otic vesicle for 30-32hpf, 36-37hpf and 48hpf, respectively. VZ: ventricular zone, region where most of progenitor cells are located. MZ: mantle zone, region of ongoing neurogenesis. Within the VZ there are dorsal progenitors (DP), medial progenitors (MP) and ventral progenitors (VP). **(E)** Representative example of transversal view at 36-37hpf, from triple whole mount smiFISH labelling neurog1 as ventral progenitor marker **(VP, yellow)**, ascl1 as medial progenitor marker **(MP, magenta)** and atoh1 as dorsal progenitor marker **(DP, cyan**). The merge image shows the three progenitor markers on their respective colors which are expressed in the ventricular zone as described on schematic in **(D)**. Dashed line indicates boundary between different progenitor regions (dorsal, medial and ventral progenitor region). Scale bar 20μm. **(F)** Representative example of transversal view at 36-37hpf, from triple whole mount smiFISH labelling pri-mir-9-5 **(cyan**) and the zebrafish mHes1 orthologues, her6 **(yellow)** and her9 **(magenta)**. The merge image shows pri-mir-9-5 **(cyan)** co-expressing with her6 **(yellow)** and *her9* **(magenta)**. Dashed line indicates boundary between different progenitor regions (dorsal, medial and ventral progenitor region). Scale bar 20μm.

To explore the identity of the triple pri-mir-9 expressing progenitors, we turn our attention to the Dorso-Ventral (D-V) progenitor axis of the ventricular zone. The everted structure of the zebrafish hindbrain means that dorsal progenitors are located more laterally than medial or ventral ones (**Fig. 4D; schematic**). We compared the expression to neurog1, ascl1 and atoh1 which are markers for ventral, medial and dorsal progenitors, respectively (**Fig. 4E**) (Tambalo et al., 2020). Remarkably, at early stages of development the cells expressing two primary transcripts were mostly localized in the ventral progenitor region of the ventricular zone (**Fig. 4A**) while at later stages the cells with three primary transcripts exclude the ventral most domain (**Fig. 4B, C**), suggesting that miR-9 high levels are required in medial and dorsal progenitors. Pri-mir-9-5 was expressed throughout the everted dorsoventral axis (**Fig. 4B, D, F**) and was co-expressed with *her6* and *her9*, both of which are expressed in the progenitor domain (mainly medial and some dorsal) and are downregulated as cells differentiate. Both contain miR-9 binding sites and are candidates for dynamic regulation by miR-9 (**Fig. 4F**) (Coolen et al., 2013; Leucht et al., 2008; Soto et al., 2020).

### Knocking out the late pri-mir-1 preferentially affects neuronal differentiation from medial progenitors

The spatial analysis above showed that the expression of pri-mir-9-1 is added onto to pre-existing pri-mir-9-4 and 9-5 expression in the medial and dorsal progenitors. To find out whether there is any specificity in deleting pri-mir-9-1 we designed a CRISPR/Cas9 based knock-down with guides that were specific to pri-mir-9-1 (**Fig. 5A, Fig. S4A-C**). This resulted in reduction on mature miR-9 and pri-mir-9-1 from 37hpf onwards when the endogenous locus is transcribed (**Fig. 5B-C**). RT-qPCR was also performed to pri-mir--9-3, 9-4 and 9-5 under mutation of pri-mir-9-1. Some reduction (with high variability between samples) was also observed in pri-mir-9-4 and 9-5 but it was not maintained at later stages of development **(Fig. S4E-F, 48hpf)**. Pri-mir-9-3 was not affected **(Fig. S4D)**.

**Figure 5.**
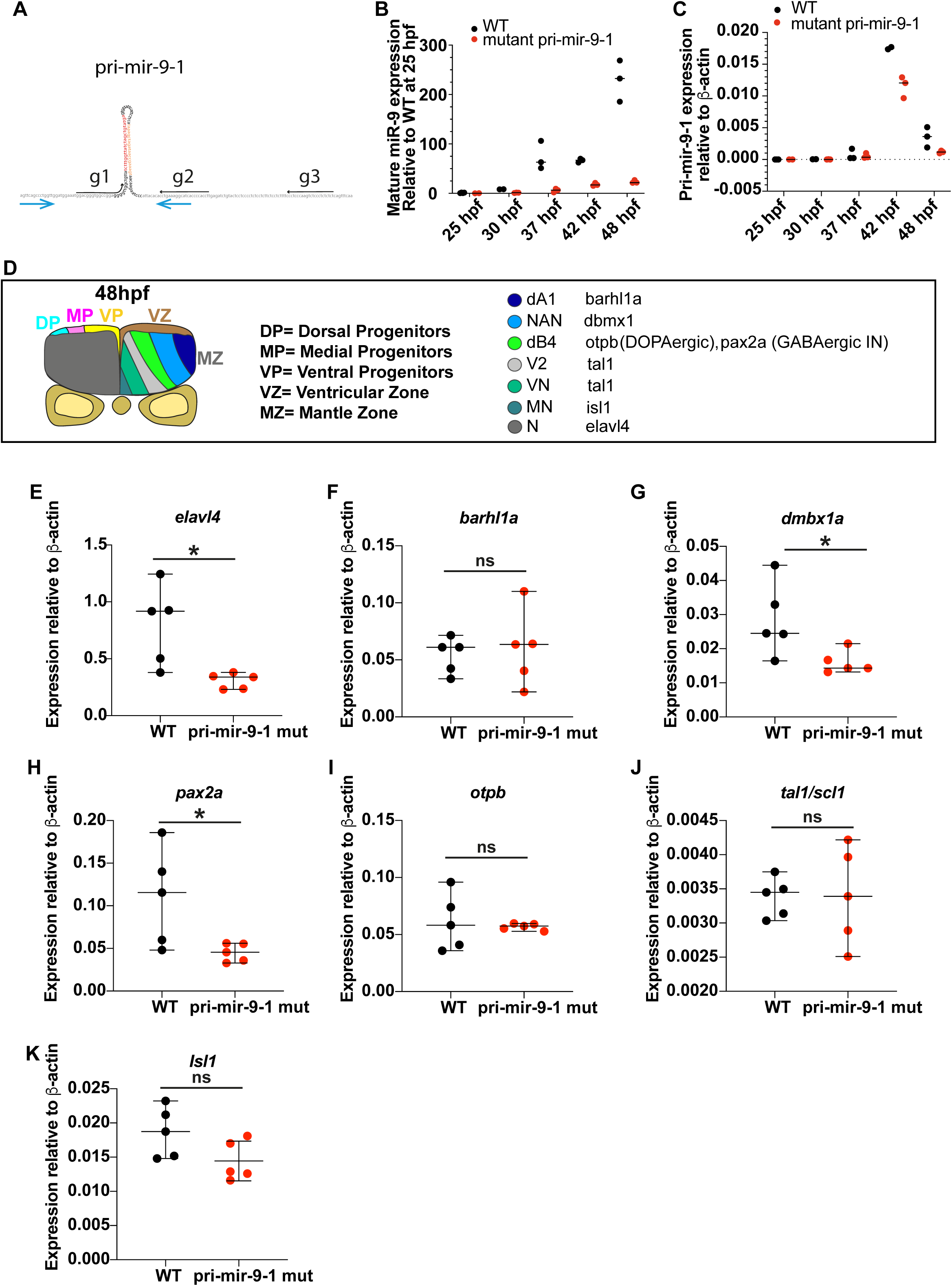
Knocking out the late pri-mir-9-1 preferentially affects neuronal differentiation from medial progenitors. **(A)** Pri-mir-9-1 hairpin loops with the respective primers used for quantitative PCR annotated as blue arrows (Methods. mRNA extraction and Quantitative PCR; Table S3). Customized guide RNA to delete specifically pri-mir-9-1 are annotated as g1, g2 and g3. Red sequence: miR-9-5’arm, orange sequence: miR-9-3’ arm, black letters: pre-mir-9, grey sequence: partial sequence of pri-mir-9-1. **(B)** Taqman RT-qPCR of mature miR-9 from dissected hindbrain at different stages of development, in wild-type conditions **(black dots)** and deletion of pri-mir-9-1 **(red dots)**, relative to wild-type at 25hpf. **(C)** SYBR green RT-qPCR relative quantification of pri-mir-9-1 from dissected hindbrain at different stages of development, in wild-type conditions **(black dots)** and deletion of pri-mir-9-1 **(red dots)**, quantification was normalised using β-actin. (**D)** Schematic representation of transverse section from zebrafish hindbrain at the level of the otic vesicle for 48h. VZ: ventricular zone, region where most of progenitor cells are located. MZ: mantle zone, region of ongoing neurogenesis. Within the VZ there are dorsal pro-genitors (DP), medial progenitors (MP) and ventral progenitors (VP). The schematic shows late neuronal markers expressed in different neuronal cell types in the hind-brain: dA1, dorsal neurons expressing *barhl1a*; NAN, noradrenergic neurons ex-pressing *dbmx1*; dB4, GABAergic interneurons expressing *pax2a*; V2, interneurons expressing *tal1*; VN, ventral neurons expressing *tal1*; MN, motor neurons expressing *isl1*; N, pan neuronal expressing *elavl4*; *otpb* is localised in dB4 region but is a marker for dopaminergic neurons. **(E-K)** SYBR green relative quantification of *elavl4* **(E)**, *barhl1a* **(F)**, *dmbx1a* **(G)**, *pax2a* **(H)**, *otpb* **(I)**, *tal1/scl1* **(J)** and *isl1* **(K)** from dissected hindbrain at 72hpf, in wild-type conditions **(black dots)** and deletion of pri-mir-9-1 **(red dots)**, quantification was normalised using β-actin. Bars indicate median with 95% confidence intervals, Mann-Whitney two-tailed test, significance p<0.05*. **(B-C)** N=3, each N contain a pool of 10 hindbrain. **(E-K)** N=5

Exploring the potential defect further we used a panel of differentiation markers spanning the D-V axis (**Fig. 5D**). Injected fish did not show overt abnormalities; however, RT-qPCR analysis at 3 dpf showed that the differentiation marker *elavl4* was reduced (**Fig. 5E**). This analysis also showed a reduction of noradrenergic neurons (NAN) derived from medial progenitors (*dbmx1a*, **Fig. 5G**) and GABAergic interneurons (*pax2a*, **Fig. 5H**) while neuronal markers from other domains where not affected (**Fig. 5 F, I, J, K**).

Medial progenitors express *her6* and *her9* (**Fig. 4F**) which are miR-9 targets and need to be downregulated before the cells can differentiate. In addition, medial/dorsal progenitors differentiate later in vertebrate development than ventral ones (Delile et al., 2019). Therefore, our findings suggest that the late activation of pri-mir-9-1 contributes to the increase of miR-9 needed to downregulate Her6/Her9 in late neural progenitors so that they can give rise to a spatiotemporally appropriate neuronal fate.

### A miR-9 stepwise increase may be required to overcome adaptation of downstream target expression

Having shown that the increase in miR-9 in development is functionally important for differentiation, we wanted to explore whether the shape of the increase is also important. In other words, whether the way that miR-9 increases in steps can be decoded. This was motivated by the biological evidence obtained from smiFISH and RT-qPCR experiments in which we observe a stepwise sharp increase of the primary transcripts (**Fig. 6A)** and the mature miR-9 **(Fig. 1B)** over time. We use a mathematical model to ask whether a simple network of gene interactions can differentially respond to stepwise increase of miR-9 rather than a gradual increase.

**Figure 6.**
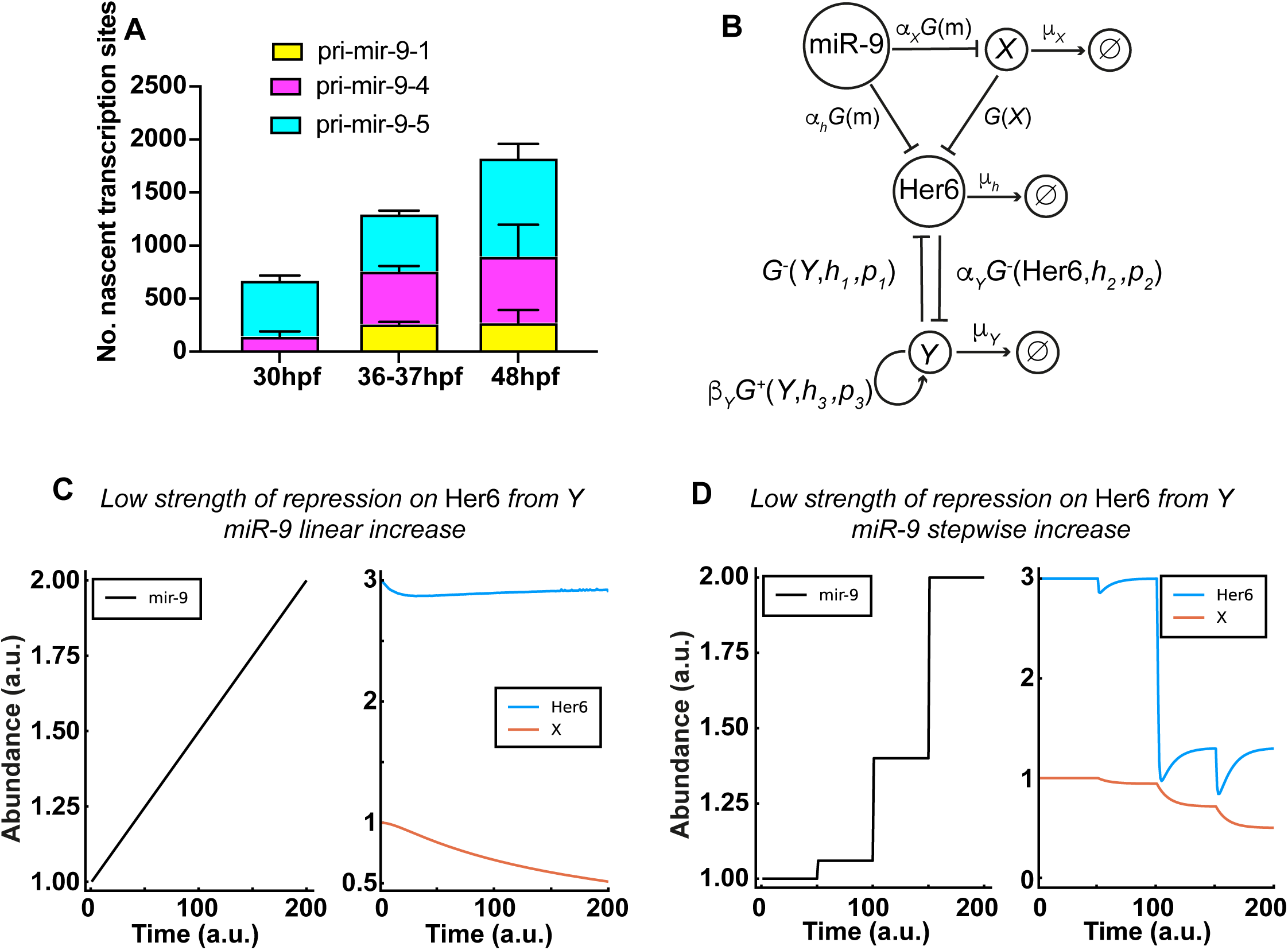
A miR-9 stepwise increase may be required to overcome adaptation of downstream target expression. **(A)** Graph representing the number of nascent transcription sites for pri-mir-9-1 **(yellow)**, pri-mir-9-4 **(magenta)** and pri-mir-9-5 **(cyan)** at 30hpf, 37-38hpf and 48hpf; in 25 microns thickness transversal section. N 30hpf= 4; N37-37hpf= 4; N 48hpf= 3. **(B)** A schematic of the extended mathematical model, which combines an incoherent feed forward loop with an additional mutually repressive, self-activating downstream target, *Y*. The parameters *α*_*h*_, *α*_*x*_ and *α*_*Y*_ represent the basal production rates of *h, X* and *Y* respectively. *µ*_*h*_, *µ*_*x*_ and *µ*_*Y*_ represent the degradation rates of *h, X* and *Y* respectively, and *β*_*Y*_ represents the production rate of Y under self-activation. The *h*_*i*_ and *p*_*i*_ are Hill coefficients and repression thresholds respectively for each of the Hill functions *G*^+^ and *G*^-^ and *G(x)* = 1/x. The model is described in detail in **Methods**, Mathematical modelling (b), and specific parameter values are listed in **Table S14. (C-D)** Dynamics of Her6 in response to different miR-9 expression profiles, for the extended model. **(C)** A linear miR-9 expression profile leads to a small initial response in Her6 expression levels, which returns to steady state levels due to the perfect adaptation. **(D)** Large instantaneous changes in miR-9 can result in a change in steady state for Her6. The initial step change is not sufficient to cause a change in steady state, therefore we introduce a fold change in the stepwise increase of miR-9, which activates *Y* and represses Her6 into a lower steady state.

Biological systems need to be robust to stochastic fluctuations that are due to low copy number, or to random perturbations in the surrounding environment. This is referred to as *adaptation* in the context of a particular output of interest, making the biological system resistant to changes of the input. However, some changes of signals are not simply due to noise or environmental fluctuations, and adaptive systems may therefore also have to respond to specific signals under changing conditions, especially during development, in order to move into a new state. Specifically, we show how a stepwise change in gene expression can allow a system to move out of an adaptively stable state. Since incoherent feed-forward loops (IFFL) are common in biology (Goentoro et al., 2009) (Shen-Orr et al., 2002) and have been shown to enable adaptation (Khammash, 2021), we hypothesized the existence of such a network centred around miR-9 as the input and Her6 as the output (**Fig. S5A; Methods**, Mathematical model) in which miR-9 affects Her6 negatively (directly) but also positively (indirectly) via repressing a repressor, *X* (IFFL). Here, miR-9 directly reduces the rate of production of Her6 protein as well as the rate of production of an intermediate (unknown) species *X*. Similarly, the production of Her6 is repressed by *X* (**Fig. S5A;** parameter values, **Table S14**). Mathematically, we say that Her6 *adapts perfectly* to changes in miR-9 since, in this model, the steady state of Her6 is independent of miR-9 (Methods Mathematical modelling (a)). The speed of this adaptation is controlled by the difference in reaction speed of the direct and indirect interactions between miR-9 and Her6. If the direct interaction is much faster than the indirect interaction, Her6 returns to steady state slowly after miR-9 copy numbers are perturbed. However, if the indirect interaction is faster, adaptation occurs quickly.

Such *perfect adaptation* is beneficial because it allows stable mean expression of Her6 in the presence of fluctuations of miR-9 expression (**Fig. S5B,C**). Conversely, no changes in miR-9, i.e linear or stepwise, can lead to persistent downregulation of Her6. Since Her6 is downregulated in response to increasing miR-9 levels during development, there would need to be an additional mechanism that enables the controlled escape from perfect adaptation. To investigate this, we extended our model to include such a potential mechanism (**Fig. 6B**). Specifically, we introduce a downstream target of Her6, named *Y*, which self-activates and interacts with Her6 through mutual repression (as we have already previously hypothesised in (Soto et al., 2020). The different behaviours of this system can be seen in **Fig. 6C-D**. A linear increase in miR-9 leads to an initial repression of Her6, which then proceeds to return to its unperturbed steady state, due to the perfect adaptation (**Fig. 6C, Fig. S6A**). However, following a sharp increase of miR-9, the concentration of Her6 decreases more strongly. This is sufficient for *Y* to overcome the repression from Her6, so that it can self-activate and in turn repress Her6 into a new, lower steady state (**Fig. 6D, Fig. S6B**). Hence, this extended motif can indeed overcome the built-in adaptation. Importantly, the escape from adaptation is triggered by a step-like change in miR-9 expression and cannot be achieved through gradual changes in miR-9 expression, or small-scale fluctuations.

The qualitative behaviour of the model is not sensitive to different values of the parameter *p*_1_, which regulates the strength of repression of Her6 by *Y*. The precise choice of *p*_1_ simply modulates the level of the lower state of Her6 expression **(Fig. 6D vs Fig. S6B)**.

## Discussion

miR-9 is expressed from several genomic loci which, after transcription and processing, produce the same 5’ mature form of miR-9 which targets the key neural progenitors transcription factors, Her/Hes. How common is this multi-locus organisation? In humans only 6.3% of mature microRNA arms are identical across two or more loci (Kozomara et al., 2019), thus, it is not very common but it is not unique to miR-9. In zebrafish this number rises to around 32.3% (Kozomara et al., 2019). The higher number of microRNA expressed from multiple loci is possibly due to the teleost-specific whole-genome duplication (WGD). Evidence from rainbow trout also shows that following the salmonid-specific extra round of WGD, microRNAs appear to be retained at higher levels than protein-coding genes (Berthelot et al., 2014). This may suggest that extra copies of microRNA are evolutionary advantageous. Here, we propose that retention of multiple microRNA loci could have specific functional advantages for regulatory control of target gene expression of an organism. By examining in detail, the temporal and spatial expression at single cell level of 3 selected early and late pri-mir-9s, from across their phylogenetic tree, we offer two possible, not-mutually exclusive, explanations for this multi-site organisation or primary transcripts.

The first explanation involves a qualitative mechanism. In this scenario, distinct pri-mir-9s have different spatial expression, which allows them to target different, i.e., region-specific, gene expression. Some differences in the spatial expression of pri-mir-9s are easily discernible at low resolution (e.g., differential expression in the forebrain) while others are subtle and require post-hybridisation sectioning to document, as we have done here. An example of the latter is the expression of pri-mir-9-1 which extents more dorsally in the hindbrain than pri-mir-9-4 at a late stage of development. This correlates well with the expression of Her6 and Her9, which are both miR-9 targets but are expressed adjacent to each other along the D-V axis (Soto et al., 2020).

The second explanation favours a quantitative mechanism. In this scenario, the differential temporal expression, where some primary transcripts commence their expression early while others are only expressed late, results in the simultaneous expression of both (or more) transcriptional loci in the same cells at a particular time in development. In support of this scenario, we have shown by smiFISH that pri-mir-9-1, a late onset primary transcript, is co-expressed in the same cells as the earlier on-set pri-mir-9-4 or 9-5. This co-expression may be a strategy to increase the amount of miR-9 available to the cell more than what would be possible with transcription from one locus alone.

Why would an increase in mature miR-9 over time be needed? One possibility is raised by the recent work from (Amin et al., 2021) who demonstrated that miRNA-dependent phenotypes emerge at particular dose ranges because of hidden regulatory inflection points of their underlying gene networks. This indicates that the miRNA cellular dose is a major determinant of *in vivo* neuronal mRNA target selection. A complementary scenario is supported by our previous work where we have shown that the dynamical profile of Hes1 (i.e. oscillatory expression to stable expression of different levels) as well as the amount of time that Hes1 oscillates for, depends on the amount of miR-9 in the cell (Goodfellow et al., 2014) (Phillips et al., 2016) (Bonev et al., 2012). More recently, we have also shown by in vivo manipulations, that the input of miR-9 changes the dynamic expression of her6 from noisy to oscillatory and then to decreasing (Soto et al., 2020).

Taken together, these findings suggest that variations in the dose level of a single miRNA achieved by additive transcription can exert regulatory effects either by targeting different downstream gene products or by modifying the dynamic expression of the same targets. Both scenarios are compatible with experimental results whereby mutating the late onset pri-mir-9-1 preferentially reduced the appearance of markers for neurons that differentiate late. This suggests that the late miR-9 increase is important for late cell fate choices.

In other cases where multiple paralogues of a microRNA have been described, differential and non-mutually exclusive qualitative and quantitative regulation may also take place. For example, a recent study found that miR-196 paralogues show both unique and overlapping expression (Wong et al., 2015). In this study, single KOs showed some unique phenotypes (qualitative mechanisms) but combinatorial KOs showed better penetrance and additional defects, suggesting an additive role of miR-196 paralogues in establishing vertebral number (quantitative mechanism).

A salient finding from our analysis is that the increase in the amount of miR-9 present in the cell is sharp, as one would perhaps expect by the onset of transcription from additional loci. An exciting possibility, supported by our mathematical modelling, is the existence of gene network motifs that do not respond to slow increases of miR-9 because they are designed to show *adaptation*, that is, to have steady output in spite of external perturbations. Such network motifs often involve IFFLs which in turn are very common in biological systems because of their multiple advantages, including fold-change detection and robustness of output (Goentoro et al., 2009) (Khammash, 2021). However, in development, cells also need to transition from one state to another in order to diversify cell fates which is essential for the development of multicellular organisms. Thus, despite the usefulness of *adaptation* for robustness and homeostasis (Khammash, 2021), a mechanism must exist to be able to over-ride it. We suggest that in the case of miR-9, a sharp, non-linear, increase may be needed to push a dynamical system into a new state and this may be associated with a cell fate change. In our case, we suggest that the increase of miR-9 during development serves to drive the dynamics of Her6 (and other targets) from one state to another, which in turn is important for the sequential acquisition of cell fates.

At present, our computational model is qualitative, rather than quantitative, and the identity of some interacting genes in the network motif are not known. Nevertheless, this model was conceptually useful to illustrate the existence of a system that can decode and distinguish between specific upstream signalling profiles. Interestingly, microRNAs are very commonly involved in transcription factor network motifs (Minchington et al., 2020), including IFFLs (Tsang et al., 2007). A fully parameterized model based on experimental evidence and identification of the unknown components/genes would be needed before it can be tested further.

In conclusion, by providing evidence for both a quantitative and qualitative mechanism, we have shed light on the possible roles of organising pri-mir-9s in several distinct genomic loci, which may have led to their evolutionary conservation. An added benefit of our work is that the detailed characterisation we have described here will enable the selection of the correct genomic locus for genetic manipulation of miR-9 production, depending on the precise spatio-temporal expression.

## Acknowledgements

We are grateful to Dr Tom Pettini for the advice on smiFISH technique, Dr Laure Bally-Cuif for sharing plasmids. The authors would also like to thank the Biological Services Facility, Bioimaging and Systems Microscopy Facilities of the University of Manchester for technical support. This work was supported by a Wellcome Trust Senior Research Fellowship to NP (090868/Z/09/Z). The funders had no role in study design, data collection and analysis, decision to publish, or preparation of the manuscript.

## Author Contributions

Conceptualization, X.S. and N.P.; Methodology, X.S. and N.P; Software, J.B., T.M. and X.S.; Validation, X.S., T.M, C.M. and R. L.; Formal Analysis, X.S. C.M., J.B. and T. M.; Investigation, X.S., R.L, J. L., C.M. and T.M.; Resources, N.P. and X.S.; Data Curation, X.S. and T.M..; Writing - Original Draft, X.S. and N.P.; Writing – Review & Editing, N.P., X.S., J.B., J.K., C.M., T.M. and M.R.; Visualization, X.S., T.M and R.L.; Supervision, N.P. and X.S; Project Administration, N.P and X.S.; Funding Acquisition, N.P.

## Methods

### Research Animals

Animal experiments were performed under UK Home Office project licenses (PFDA14F2D) within the conditions of the Animal (Scientific Procedures) Act 1986. Animals were only handled by personal license holders.

### mRNA extraction and Quantitative real-time PCR (RT-qPCR)

miRNAs and total mRNA were extracted from a pool of 10 zebrafish hindbrains using the miRVana miRNA Isolation kit and gDNA removed using DNase1 (NEB). Reverse transcription was performed with either TaqMan MicroRNA Reverse Transcription kit (Applied Biosystems) for mature miR-9 or SuperScript III (Invitrogen) with random hexamers for pri-miRNAs. Each qPCR reaction was prepared in triplicate in a 96-well plate with the relevant TaqMan MicroRNA assay or using POWER SYBR Green Mastermix (ThermoFisher Scientific), 0.2 μM each forward and reverse primer (See **Table S3** for respective primers). and 50ng cDNA. Reactions were run on Step One Plus Real-time PCR System (Applied Biosystems) alongside negative controls The data for each sample was normalized to the expression level of U6 snRNA for mature miR-9 or *b-actin* for pri-miR9s and analysed by the 2^−ΔΔCt^ method.

For each primer pair, the PCR product was examined by gel electrophoresis and its melting curve to ensure a single fragment of the predicted molecular weight.

### Molecular cloning

RNA probes for pri-mir-9-1, pri-mir-9-2, pri-mir-9-4, pri-mir-9-5 and pri-mir-9-7 were PCR amplified and cloned into pCRII vector using primers described in **Table S1**. Except for pri-mir-9-2 probe, they were designed to distinguish the primary transcripts by including sequences, intron and exon, before and after each microRNA processing, while also covering the sequence corresponding to mature miR-9 **(Figure S2)**. Since the mature miR-9 sequence is conserved between paralogs, to avoid any cross-binding of probes to this sequence we mutated it on each probe by using QuikChange II XL Site-Directed Mutagenesis assay. This allowed us introduce deletions and single nucleotide exchange in specific regions of the mature miR-9 sequence (**Table S2**; **Figure S2**; sequence highlighted in red).

pri-mir-9-3 and pri-mir-9-6 probes were generated from plasmids kindly gifted by Laure Bally-Cuif (Nepal et al., 2016).

### Whole mount chromogenic and fluorescence *in situ* hybridization and sectioning

Chromogenic *in situ* hybridisation was performed as previously described by Christine Thisse (Thisse and Thisse, 2008). Multicolour fluorescence *in situ* hybridisation was modified from Hoppler and Vize (Lea et al., 2012) by developing with tyramide amplification (Perkin Elmer) after addition of antisense RNA probes and antibodies conjugated to horseradish peroxidase (Lea et al., 2012).

Transveral sections were obtained as described in Dubaissi (Dubaissi et al., 2012) with modifications. Embryos were embedded in 25% fish gelatine and 30% sucrose for a minimum of 24 hrs. 18μm thickness hindbrain sections were collected and transferred onto superfrost glass slides. The slides were air dried overnight under the fume hood and mounted with Prolong Diamond Antifade.

### Imaging

Chromogenic *in situs* were imaged using a Leica M165FC with a DFC7000T camera. Fluorescent *in situ* sections were imaged using Leica TCS SP5 upright confocal with HCX PL APO LU-V-I 20×0.5 water UV lens or Olympus FLUOVIEW FV1000 confocal with UPLSAPO 20X NA:0.75 lens.

### smiFISH probe design and synthesis

The smiFISH probes were designed using the probe design tool at http://www.biosearchtech.com/stellarisdesigner/. The software can assign varied size of probes, 18-22 nt, therefore we gave a size of 20nt for all designed probes with the maximum masking level available for zebrafish. Using the respective pri-mir-9 sequence we designed 36 probes for pri-mir-9-1, 35 probes for pri-mir-9-4 and 35 probes for pri-mir-9-5. Using the respective gene mature mRNA sequence, we designed 29 probes for *her6*, 33 probes for *her9*, 40 probes for *neurog1*, 39 probes for *atoh1a* and 40 probes for *ascl1a* (**Table S5-S12, respectively**). The designed probes were X-FLAP tagged (5’CCTCCTAAGTTTCGAGCTGGACTCAGTG3’) at the 5’ of each gene-specific sequence. The gene specific probes were ordered from IDT in a 96 well format in nuclease-free water, 100μM concentration. Upon arrival, we combined 100 μl of the gene-specific probe together, mixed, split into 100 μl aliquots and stored at -20°C. In addition, we ordered fluo-FLAP sequences (5’CACTGAGTCCAGCTCGAAACTTAGGAGG3’) from either IDT or Biosearch Technology, they were labelled with either Atto-550, CalFluor-610 or AlexaFluor-647. Each gene specific probe mix was labelled by mixing 2μl of the gene-specific X-FLAP probe mix (100μM), 2.5μl of fluo-FLAP (100μM) and 5μl of 10x NEB buffer 3 in a final volume of 50μl. The hybridisation cycle was 85°C 3 min, 65°C 3 min and 25°C 3 min. The labelled probe was stored at -20°C.

### Whole mount smiFISH

Whole-mount smiFISH protocol for zebrafish embryos was developed by adapting smiFISH protocol from (Marra et al., 2019). Embryos were fixed in 4% formaldehyde in 1x PBS. After smiFISH protocol, embryos were stained with Phalloidin-Alexa Fluor 488 (400x dilution in PBS 1x-Tween 0.1%) for 1h at room temperature and followed by 3 washes with PBS-Tween. Embryos were embedded in 4% low melting point agarose (Sigma) to collect 250μm thickness hindbrain transversal sections.

### smiFISH microscopy and deconvolution

smiFISH images were collected using Leica TCS SP8 upright confocal with HC APO L U-V-I 63x/0.9 water lens, magnification 0.75x. We acquired three-dimensional stacks 1024 × 1024 pixels and z size 0.63μm. magnification 0.75x. Bits Per Pixel: 16. Pinhole: 1 Airy Unit. Scan speed: 200. Channels were sequentially imaged. smiFISH images were collected with frame accuracy 3 and line average 6.

Deconvolution of confocal images was performed using Huygens Professional Software. As pre-processing steps, the images were adjusted for (1) ‘Microscopic Parameters’ and for (2) ‘object stabilizer’ as additional restoration, the latter was used to adjust for any drift during imaging. Following, we used the deconvolution Wizard tool, the two main factors to adjust during deconvolution were the background values and the signal to noise ratio. Background was manually measured for every image and channel, while the optimal signal to noise ratio identified for the images was value 3. After deconvolution the images were generated using Imaris 9.5.

### smiFISH segmentation

Segmentation was performed using Phalloidin-AlexaFluor 488 as membrane marker. Using imaris 9.5 software we selected the ‘Cells tool’ from which ‘Cells only’ was used as detection type and ‘Cell boundary’ was selected as cell detection type. Automated segmentation was performed followed by manual curation to identify for cells incorrectly segmented.

To quantify pri-mir-9 nascent transcriptional sites we used the ‘Spot tool’. The estimated spot diameter size was XY 1μm and Z 2μm. We used the default parameters to identify the nascent transcriptional sites and further manual curation was performed to correct for minimal errors carried out by the software. Further on, spots were imported into the segmented cells to identify the cells that contained one, two or three pri-mir-9s.

### Expression analysis of hes/her genes and microRNA hosts

For the *in silico* analysis of the microRNA host gene expression we downloaded the time course RNA-seq data (TPM) from (White et al., 2017) supplemental file 3. Here we used the overlapping host genes as a proxy for the expression of the microRNA. MicroRNA would not show up in standard RNA-seq analysis and there is no current microRNA time course data. Host genes were identified as those with overlapping annotations with the miR-9 genes. The host genes for each pri-mir-9 are in **Table S4**. Pri-mir-9-7 has no overlapping annotation at this time and is thus not reported on in these data.

We filtered the RNA-seq data removing genes which were neither the host genes of the microRNA or members of the Her family. 3 repeats for each stage of development are included in the data and we averaged the expression across the 3 repeats for each stage. The stages reported in the data are based on standard embryonic stages in zebrafish development. However, we wanted to visualize the expression in terms of hours and the stages were converted accordingly. Finally, before plotting these data were z-scored to normalize the expression of each of the genes so that we could compare changes in expression over time rather than absolute levels. These data were then plotted using the heatmap.3 package in R.

### Deletion of pre-mir-9-1 using CRISPR/Cas9

For pre-mir-9-1 deletion using CRISPR/Cas9, sgRNA target sites were identified using the CRISPRdirect (http://crispr.dbcls.jp/) and Target Finder (Feng Zhang lab http://crispr.mit.edu/). sgRNAs were generated following CRISPRscan protocol (Moreno-Mateos, Vejnar et al., 2015)) using the oligonucleotides described in **Table S13**. Transcription of sgRNA was carried out using MEGAshortscript T7 kit (Ambion/Invitrogen) with 100-400ng of purified DNA following the manufacturer’s instructions. After transcription sgRNA was purified using MEGAclear™ Transcription Clean-Up Kit. The Cas9nls protein was obtained from NEB (cat number M0646T).

One-cell stage wild type embryos were injected with ∼1 nl of a solution containing 185 ng/μl Cas9nls protein, 125 ng/μl sgRNA, 40 ng/μl caax-mRFP mRNA in 0.05% phenol red. To evaluate if each sgRNA was generating mutation, genomic DNA was extracted from 3-4dpf embryos using 50 μl NP lysis buffer per embryo (10mM Tris pH8, 1 mM EDTA, 80mM KCl, 0.3 % NP40 and 0.3% Tween) and 0.5 μg/μl Proteinase K (Roche) for 3-4 hrs at 55°C, 15 min at 95°C and then stored at 4°C. Then, High Resolution Melt (HRM) was performed **(Fig. S4A)** using Melt Doc kit (Applied Biosystems) following manufacturer instructions, specific primers were designed to generate an amplicon of 395bp in wildtype conditions: forward primer 5’ACAGTTGACTTTCTAATTACAACCC-3’ and reverse primer 5’AGCAGGAGGAGATAATCACAGC-3’.

To generate pre-mir-9 deletion we combined three different sgRNA flanking the region of the mature miR-9 region, they were microinjected as described above. We chose using three sgRNA to increase our probability deleting the mature miR-9 sequence. Embryos were injected with minimal amounts of sgRNA (F0) to not have overt phenotype at the macroscopic level during the experimental period (24hpf-72hpf) thus, minimizing the chances of non-specific toxicity. Further on, the amplicons with deletion were identified by agarose gel and sequenced (**Fig S4B,C)**.

### Mathematical modelling

#### (a) Steady state calculation of Her6 in the perfect adaptation model

The perfect adaptation model can be described by a set of differential equations (**Fig. S5A and Table S14**):

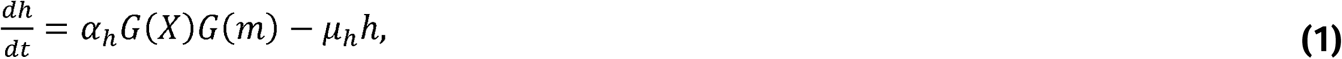

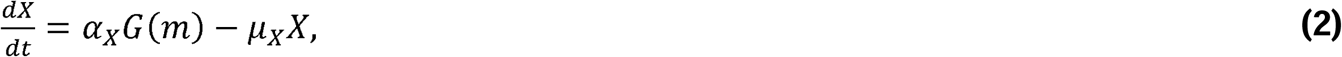

where *h* is Her6, *m* is miR-9, and *α*_*h*_, *µ*_*h*_, *α*_*x*_ and *µ*_*x*_ are positive real constants which represent the production and degradation rates of Her6 and *X* respectively. The negative interaction between each of these model components is given by an arbitrary function G. To identify a possible shape of G, we consider the steady state of eqs (1) and (2), which leads to:

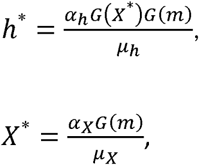

which combine to give

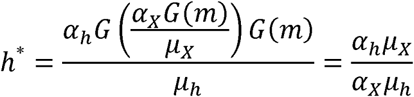

if *G* is defined by the negative interaction, *G*(*m*) = 1/*m*. Hence, in our chosen model the steady state of Her6, *h*\*, is independent of miR-9. To achieve this, we made the assumption that G is a nonlinear negative interaction, which agrees with previous models of miR-9 interactions (Goodfellow et al., 2014). In order to explore the adaptation properties of this network, we made certain simplifications over previous models (Goodfellow et al., 2014) such as omitting the her6 autorepression, transciptional delays and noise. Thus, this simplified Her6 network does not reproduce the oscillatory expression of Her6 but instead explores the transition between different stable steady states.

#### (b) Extended model

The extended system can be described by the following set of differential equations **(Fig. 6A and Table S14)**:

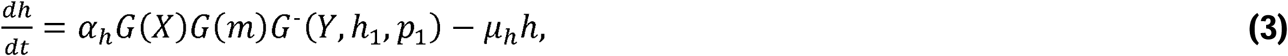

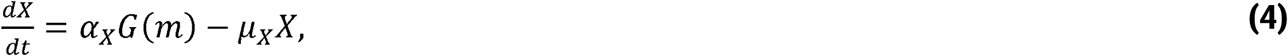

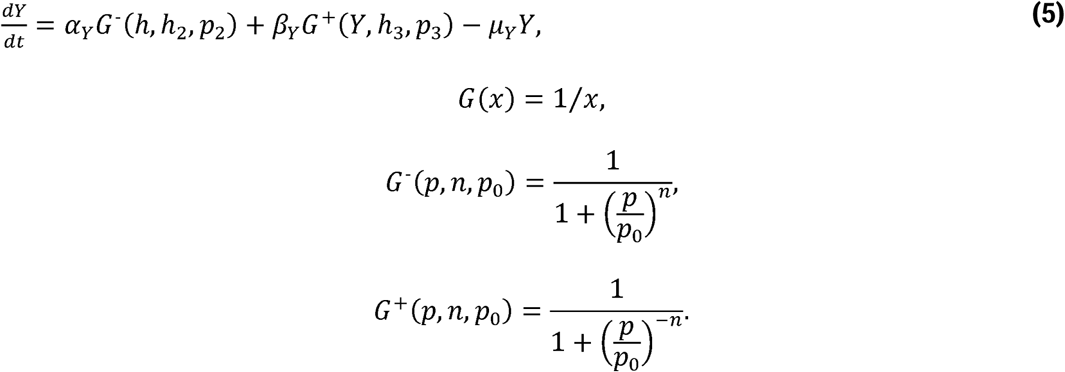

We pre-define the profile of miR-9 expression over time to interrogate both stepwise and linear expression, and then solve the system for *h, X* and *Y*. The parameters *α*_*h*_, *α*_*x*_ and *α*_*Y*_ represent the basal production rates of *h, X* and *Y* respectively. Similarly, *µ*_*h*_, *µ*_*x*_ and *µ*_*Y*_ represent the degradation rates of *h, X* and *Y* respectively, and *β*_*Y*_ represents the production rate of Y under self-activation. The *h*_*i*_ and *p*_*i*_ are Hill coefficients and repression thresholds respectively for each of the Hill functions. For the activating Hill function *G*^+^(*p, n, p*_0_) with arbitrary input parameter *p*, Hill coefficient *n* and repression threshold *p*_0_, as *p* grows much larger than *p*_0_, *G*^+^ tends to 1, and as *p* goes to 0, *G*^+^ tends to 0. For *G*^-^ the limits are reversed, i.e. *G*^-^ is equal to 1 for small values of *p* and goes to 0 for for *p* » *p*_0_. The Hill coefficient *n* determines the sensitivity of the function to changes in *p*, i.e. larger *n* corresponds to higher sensitivity. All parameters introduced here are constants, and their values are listed in Table S14. These parameters are chosen such that *Y* is repressed and has no effect on the system when Her6 is at its high steady state *h*^*^.

