## supplemental information for "Timing of neurogenesis through sequential accumulation of miR-9 due to additive expression of multiple alleles"

**TABLE OF CONTENTS**

Figure S1 - S6

Figure Legend S1 - S6

Table S1 - S14

### Figure S1

A

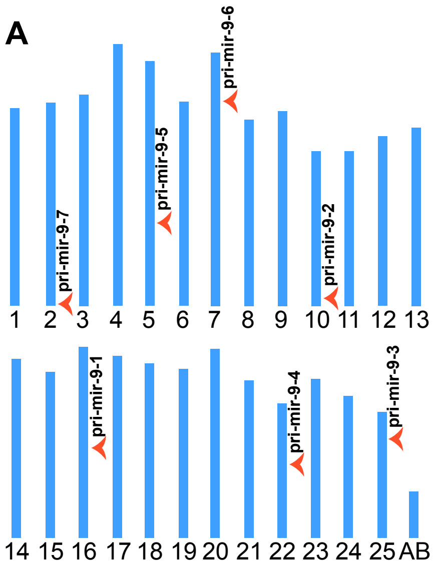


B

| name | chromosome | Genomic location | strand |
|---|---|---|---|
| Pri-mir-9-1 | chr16 | lincRNA, intragenic/exonic | + |
| Pri-mir-9-2 | chr10 | lincRNA, intragenic/ intronic | - |
| Pri-mir-9-3 | chr25 | intergenic | - |
| Pri-mir-9-4 | chr22 | BX664610.1 protein, intragenic/intronic | + |
| Pri-mir-9-5 | chr5 | lincRNA, intragenic/exonic | - |
| Pri-mir-9-6 | chr7 | lincRNA, intragenic/exonic | - |
| Pri-mir-9-7 | chr2 | lincRNA. intragenic/intronic | - |

C

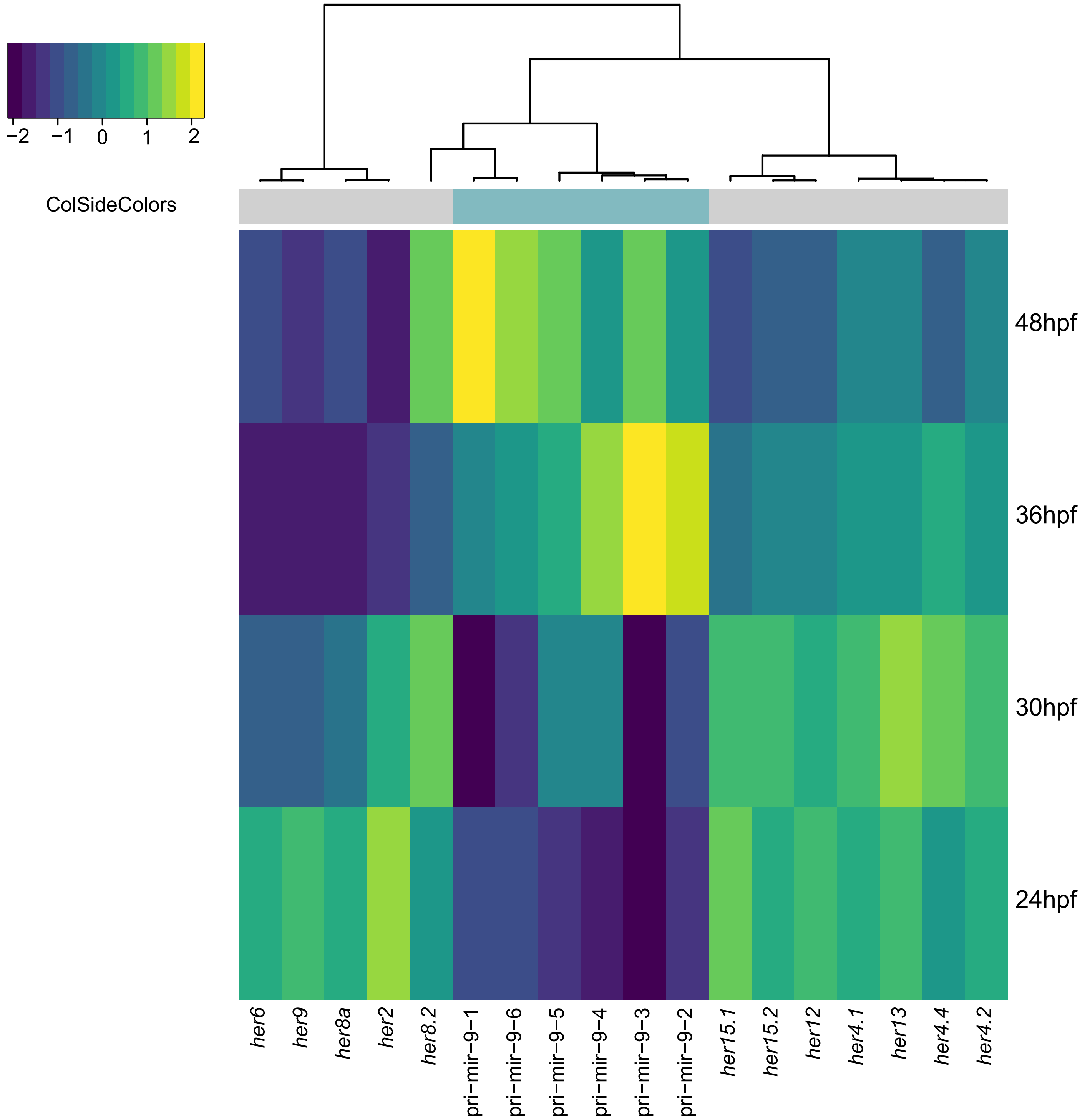

#### Figure S2

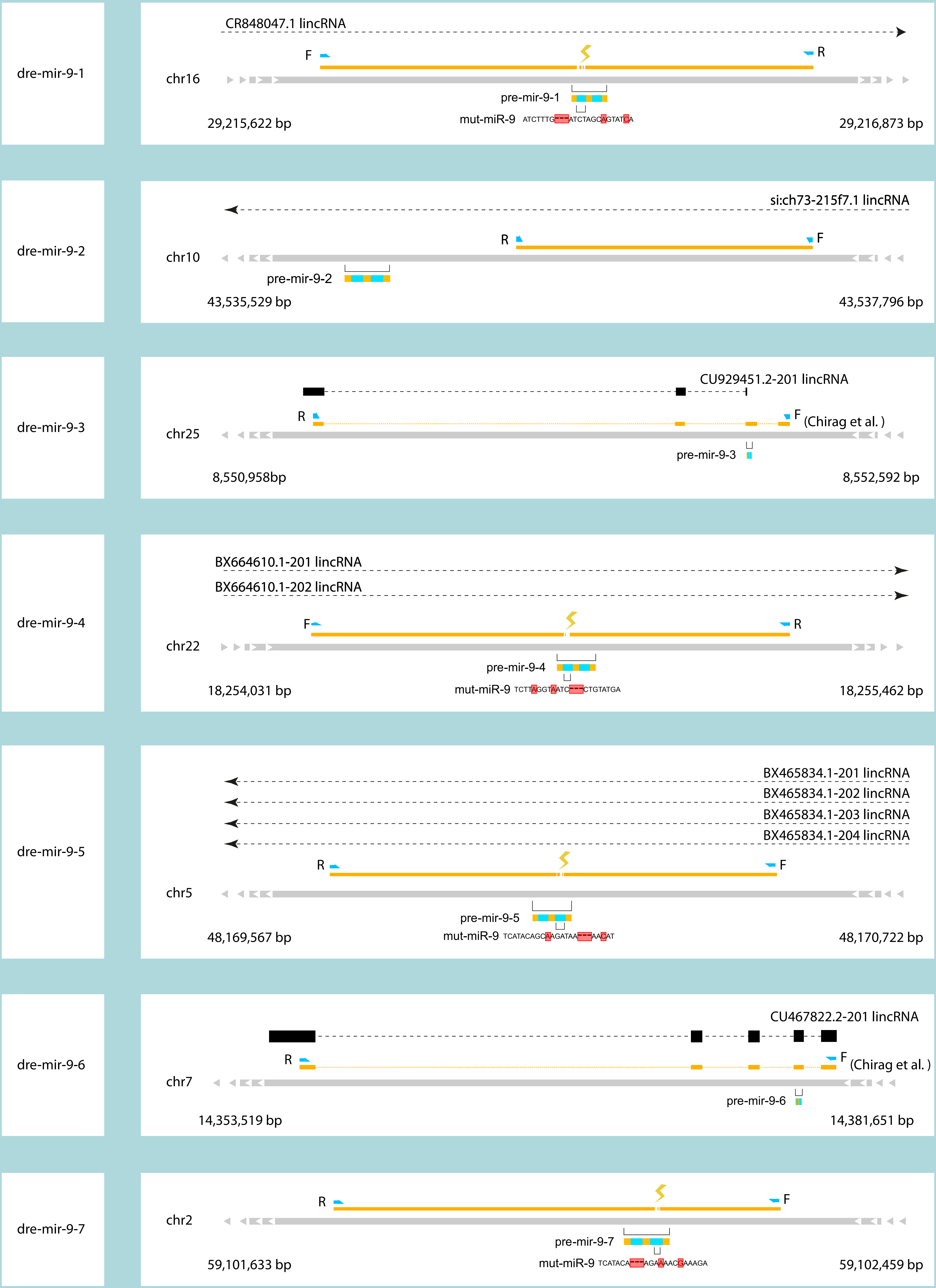
dre-mir-9-1
CR848047.1 lincRNA
F
R
chr16
pre-mir-9-1
mut-miR-9 ATCTTTG---ATCTAGCAGTATCA
29,215,622 bp
29,216,873 bp
dre-mir-9-2
si:ch73-215f7.1 lincRNA
R
F
chr10
pre-mir-9-2
43,535,529 bp
43,537,796 bp
dre-mir-9-3
CU929451.2-201 lincRNA
R
F (Chirag et al. )
chr25
pre-mir-9-3
8,550,958bp
8,552,592 bp
dre-mir-9-4
BX664610.1-201 lincRNA
BX664610.1-202 lincRNA
F
R
chr22
pre-mir-9-4
mut-miR-9 TCTTAGGTAATC---CTGTATGA
18,254,031 bp
18,255,462 bp
dre-mir-9-5
BX465834.1-201 lincRNA
BX465834.1-202 lincRNA
BX465834.1-203 lincRNA
BX465834.1-204 lincRNA
R
F
chr5
pre-mir-9-5
mut-miR-9 TCATACAGCAAGATAA---AACAT
48,169,567 bp
48,170,722 bp
dre-mir-9-6
CU467822.2-201 lincRNA
R
F (Chirag et al. )
chr7
pre-mir-9-6
14,353,519 bp
14,381,651 bp
dre-mir-9-7
R
F
chr2
pre-mir-9-7
mut-miR-9 TCATACA---AGAAAACGAAAGA
59,101,633 bp
59,102,459 bp

#### Figure S3

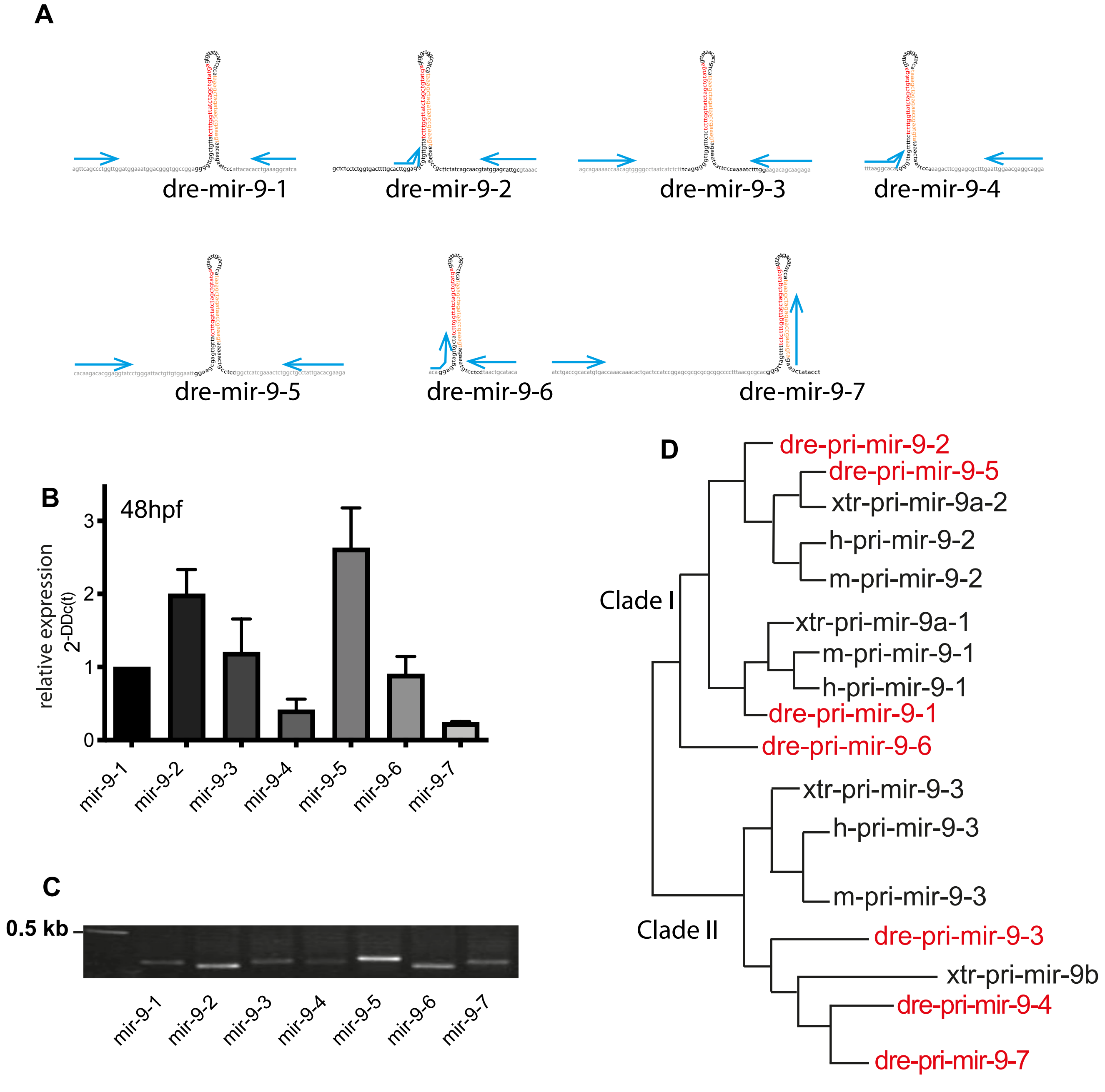

A
dre-mir-9-1
dre-mir-9-2
dre-mir-9-3
dre-mir-9-4
dre-mir-9-5
dre-mir-9-6
dre-mir-9-7
B
48hpf
relative expression
2-DDc(t)
0
1
2
3
mir-9-1
mir-9-2
mir-9-3
mir-9-4
mir-9-5
mir-9-6
mir-9-7
C
0.5 kb
mir-9-1
mir-9-2
mir-9-3
mir-9-4
mir-9-5
mir-9-6
mir-9-7
D
Clade I
Clade II
dre-pri-mir-9-2
dre-pri-mir-9-5
xtr-pri-mir-9a-2
h-pri-mir-9-2
m-pri-mir-9-2
xtr-pri-mir-9a-1
m-pri-mir-9-1
h-pri-mir-9-1
dre-pri-mir-9-1
dre-pri-mir-9-6
xtr-pri-mir-9-3
h-pri-mir-9-3
m-pri-mir-9-3
dre-pri-mir-9-3
xtr-pri-mir-9b
dre-pri-mir-9-4
dre-pri-mir-9-7

### Figure S4

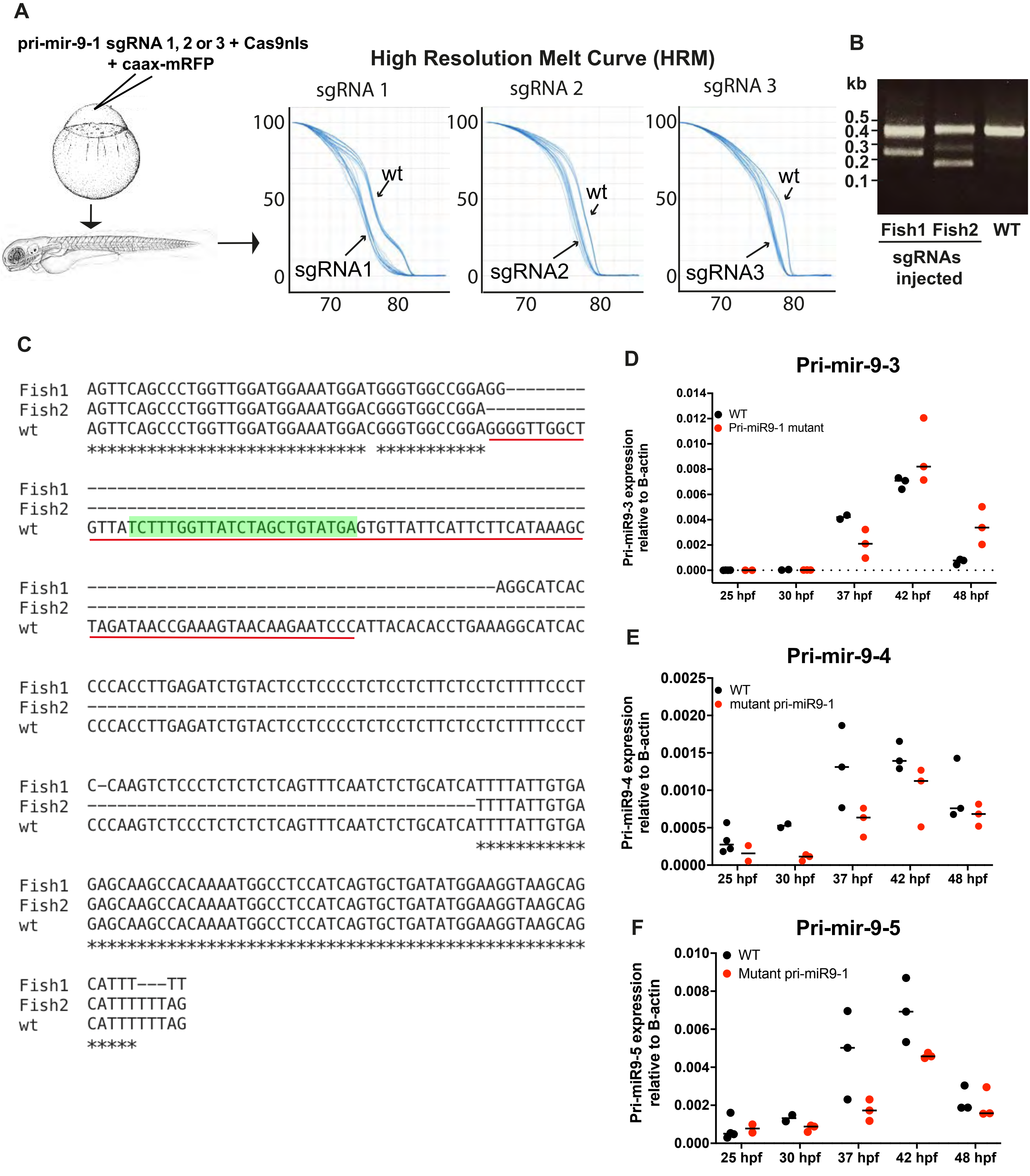

A
pri-mir-9-1 sgRNA 1, 2 or 3 + Cas9nls
+ caax-mRFP
High Resolution Melt Curve (HRM)
sgRNA 1
sgRNA 2
sgRNA 3
100
50
0
70
80
wt
sgRNA1
sgRNA2
sgRNA3
B
kb
0.5
0.4
0.3
0.2
0.1
Fish1 Fish2
WT
sgRNAs injected
C
Fish1 AGTTCAGCCCTGGTTGGATGGAAATGGATGGGTGGCCGGAGG--------
Fish2 AGTTCAGCCCTGGTTGGATGGAAATGGACGGGTGGCCGGA----------
wt AGTTCAGCCCTGGTTGGATGGAAATGGACGGGTGGCCGGAGGGGTTGGCT
**************************** ***********
Fish1 --------------------------------------------------
Fish2 --------------------------------------------------
wt GTTATCTTTGGTTATCTAGCTGTATGAGTGTTATTCATTCTTCATAAAGC
Fish1 -----------------------------------------AGGCATCAC
Fish2 --------------------------------------------------
wt TAGATAACCGAAAGTAACAAGAATCCCATTACACACCTGAAAGGCATCAC
Fish1 CCCACCTTGAGATCTGTACTCCTCCCCTCTCCTCTTCTCCTCTTTTCCCT
Fish2 --------------------------------------------------
wt CCCACCTTGAGATCTGTACTCCTCCCCTCTCCTCTTCTCCTCTTTTCCCT
Fish1 C-CAAGTCTCCCTCTCTCTCAGTTTCAATCTCTGCATCATTTTATTGTGA
Fish2 ---------------------------------------TTTTATTGTGA
wt CCCAAGTCTCCCTCTCTCTCAGTTTCAATCTCTGCATCATTTTATTGTGA
***********
Fish1 GAGCAAGCCACAAAATGGCCTCCATCAGTGCTGATATGGAAGGTAAGCAG
Fish2 GAGCAAGCCACAAAATGGCCTCCATCAGTGCTGATATGGAAGGTAAGCAG
wt GAGCAAGCCACAAAATGGCCTCCATCAGTGCTGATATGGAAGGTAAGCAG
**************************************************
Fish1 CATTT---TT
Fish2 CATTTTTTAG
wt CATTTTTTAG
*****
D
Pri-mir-9-3
Pri-miR9-3 expression relative to B-actin
0.014
0.012
0.010
0.008
0.006
0.004
0.002
0.000
WT
Pri-miR9-1 mutant
25 hpf
30 hpf
37 hpf
42 hpf
48 hpf
E
Pri-mir-9-4
Pri-miR9-4 expression relative to B-actin
0.0025
0.0020
0.0015
0.0010
0.0005
0.0000
WT
mutant pri-miR9-1
25 hpf
30 hpf
37 hpf
42 hpf
48 hpf
F
Pri-mir-9-5
Pri-miR9-5 expression relative to B-actin
0.010
0.008
0.006
0.004
0.002
0.000
WT
Mutant pri-miR9-1
25 hpf
30 hpf
37 hpf
42 hpf
48 hpf

### Figure S5

#### PERFECT ADAPTATION MODEL

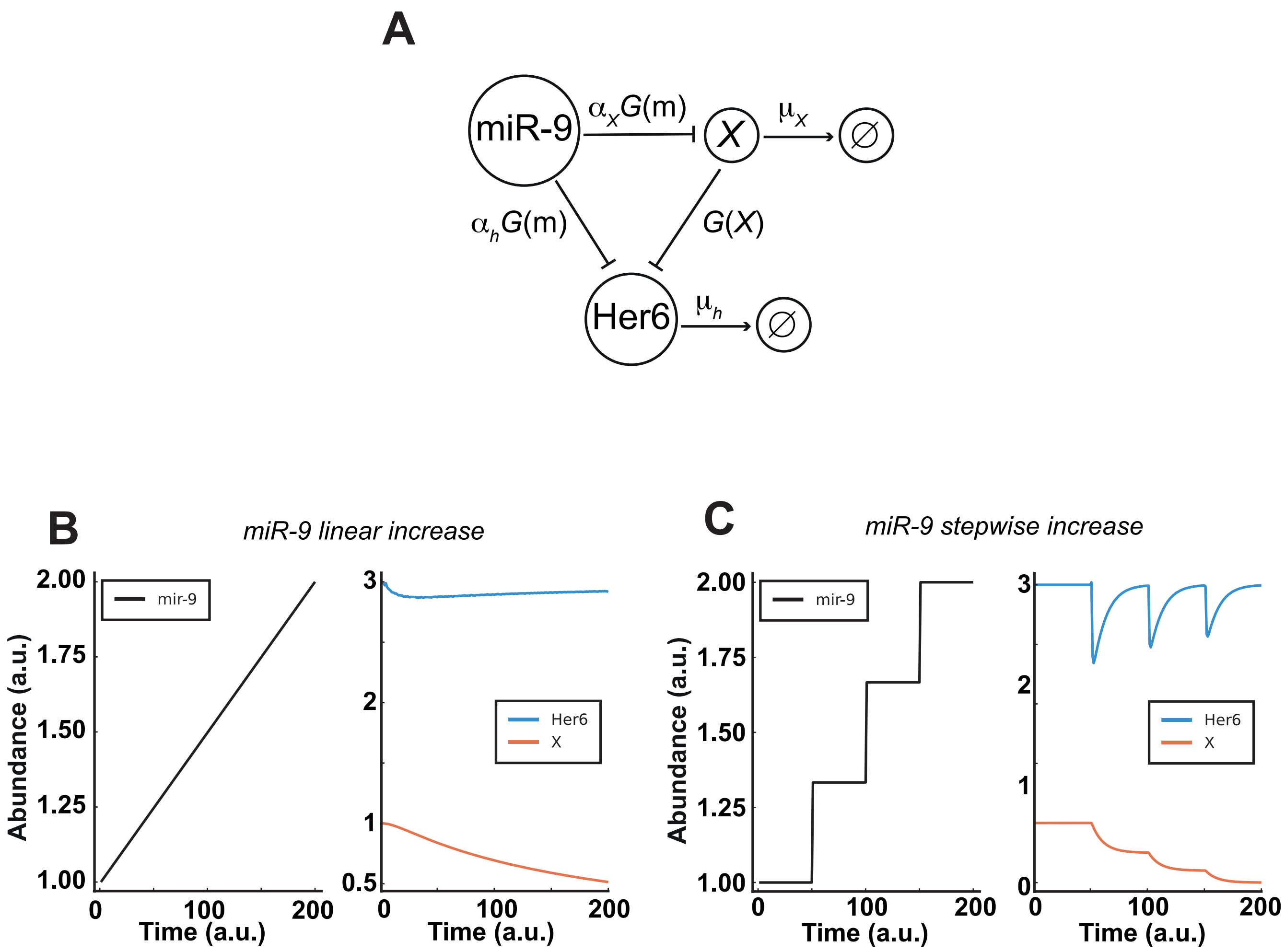

### Figure S6

#### EXTENDED MODEL

**A** *High strength of repression on* Her6 *from Y*
*miR-9 linear increase*

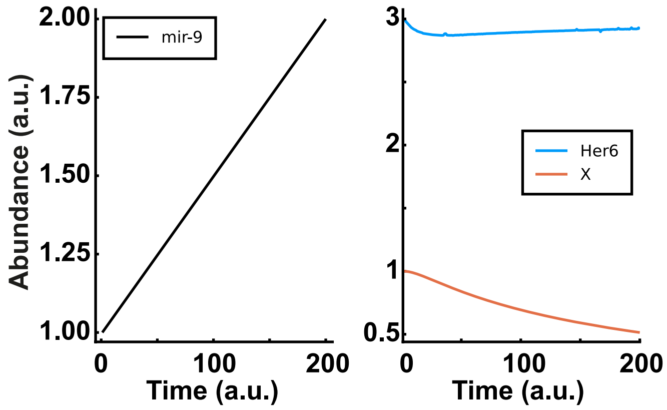


**B** *High strength of repression on* Her6 *from Y*
*miR-9 stepwise increase*

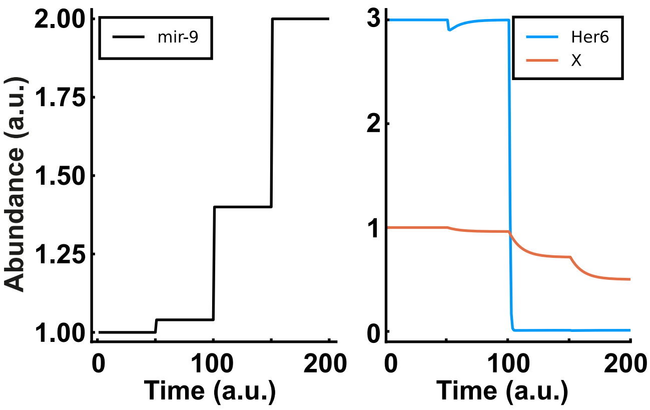

#### Supplemental Figure Legend

**Figure S1. miR-9 host gene expression negatively correlates with Hes/Her gene expression over developmental time. (A)** Map of miR-9 paralogues in Danio rerio produced with information from Vega genome browser, data was collated from ensembl genome browser GRCz10, showing chromosomal loci. Red head arrow indicates location of respective pri-mir-9. **(B)** Table showing the genomic loci of the seven Danio rerio miR-9 paralogues, their host gene, exonic or intronic location within the host gene and strand orientation**. (C)** Heatmap showing z-scored expression data over developmental time for the miR-9 host / pri-mirs and Hes/Her genes over developmental time in the zebrafish embryo. Data is from White *et al.*, 2017.

**Figure S2. Probe design for *in situ* hybridisations of pri-mir-9s.** The probes are indicated by the orange blocks. Where probes are on exons, introns are indicated by dashed lines. Forward (F) and reverse (R) are visualised by blue arrows. Overlapping transcripts are indicated by the black boxes and dashed lines. The microRNA hairpin is indicated by a small orange box broken by two blue boxes indicating the mature sequences. Probes for pri-mir-9-3 and pri-mir-9-6 were based on those from Chirag et al. Mutations in the mature sequence are indicated by lightning bolts and also annotated underneath.

**Figure S3. All seven miR-9 paralogues are expressed at 48hpf. (A)** Schematic of the seven miR-9 paralogues hairpin loops with the respective primers used for qRT-PCR annotated as blue arrows **(Methods. mRNA extraction and Quantitative real-time PCR; Table S3).** Red sequence: miR-9-5'arm, orange sequence: miR-9-3' arm, black letters: pre-mir-9, grey sequence: partial sequence of pri-mir-9. **(B)** Bar plot showing relative expression of the seven miR-9 paralogues at 48hpf, N=3. Error bar represent mean and SEM. **(C)** Amplicons of the seven miR-9 paralogues generated by qRT-PCR using the respective primers **(Table S3). (D)** Evolutionary relationship of pri-mir-9 family modified from Alwin Prem Anand et al., 2018 phylogenetic analysis. Highlighted in red letter are the seven miR-9 paralogues. The zebrafish miR-9 paralogues cluster into two Clades: Clade I that is divided into 2 subgroups: Subgroup I corresponding to the group in which human (h) and mouse (m) pri-mir-9-2 precursor cluster with danio rerio (dre) pri-mir-9-2 and pri-mir-9-5 precursors and subgroup II corresponding to the group where h/m-pri-mir-9-1 precursor clusters with dre-pri-mir-9-1. Interestingly, dre-pri-mir-9-6 is closely related to clade I but doesn't group with any subgroup. Clade II correspond to the branch where m/h-pri-miR-9-3 group with dre-pri-mir-9-3, dre-pri-mir-9-4 and dre-pri-mir-9-7.

**Figure S4. Deletion of pre-mir-9-1**

**(A). (A-left)** Schematic representation of experimental procedure used to delete pre-mir-9-1, **(A-right)** high resolution melt graphs obtained from: wt (wildtype) versus sgRNA 1, 2 or 3 injected fish, respectively. **(B)** Agarose gel showing the size of the amplicon in wt fish (395bp) and in injected with three sgRNA fish, ~250bp and 395bp in fish1 and ~190bp and 395bp in fish2. **(C)** Representative examples of sequences obtained from F0 embryos showing deletion of pre-mir-9-1. Red underline indicates pre-mir-9-1 sequence. Green highlighted indicates mature miR-9 sequence. **(D-F)** SYBR green relative quantification of pri-mir-9-3 **(D),** pri-mir-9-4 **(E)** and pri-mir-9-5 **(F)** from dissected hindbrain at different stages of development, in wild-type conditions

**(black dots)** and deletion of pre-mir-9-1 **(red dots)**, quantification was normalised using β-actin. N=3.

**Figure S5. Her6 is not down-regulated by miR-9 in the perfect adaptation model**

**(A-C) Perfect adaptation model. (A)** A schematic of the perfect adaptation model based on an incoherent feed forward loop. The parameters $\mu_h$, $\alpha_h$ and $\alpha_X$ are positive real constants. $\mu_h$ represents the degradation rates of Her6 and $\alpha_h$ and $\alpha_X$ represent the basal production rate of Her6 and $X$ respectively. See **Methods,** Mathematical modelling (a), parameter values are given in **Table S14. (B)** A linear mir-9 expression profile leads to a small initial response in Her6 expression levels, which returns to steady state levels due to the perfect adaptation. **(C)** Large instantaneous changes in miR-9 cause a large drop in Her6 levels, which then returns to steady state. The model detects fold changes in miR-9 and repetitive steps result in a diminished response from Her6.

**Figure S6. Her6 is down-regulated by miR-9 stepwise increase in extended adaptation mathematical model**

**(A-B)** Dynamics of Her6 in response to linear increase in miR-9 **(A)** or stepwise increase in miR-9 **(B)** with increased repression strength from *Y*. The stepwise increase in miR-9 under high repression strength from *Y* causes Her6 to switch off given a large enough increase in miR-9.

**Table S1.** Sequences of primers used to generate probes for *in situ* hybridisation. * Primers for *Pri-mir-9-3* and *Pri-mir-9-6* were used to clone the probes that were kindly provided by the Bally-Cuif lab.

| Gene | Oligonucleotides sequence 5>3 |
|---|---|
| Pri-mir-9-1 f | cagattgacagagttgtgag |
| Pri-mir-9-1 r | cagtgctgactactctaatg |
| Pri-mir-9-2 f | gttcaatcctcttccgtttg |
| *Pri-mir-9-2 r | cagcatcccgttacacattc |
| *Pri-mir-9-3 f | aatgctgagttttgccacct |
| Pri-mir-9-3 r | tgctgcggaaaataacacaa |
| Pri-mir-9-4 f | ttccacaagggtatcgatag |
| Pri-mir-9-4 r | tattatatgagaaccacgtg |
| Pri-mir-9-5 f | gatgttatttccgcgtgcac |
| Pri-mir-9-5 r | tcagctctctctatatgtcc |
| *Pri-mir-9-6 f | gagagacaacatcgcatcca |
| *Pri-mir-9-6 r | tcaaacatagcaggatagggtct |
| Pri-mir-9-7 f | tactaattaacctataacgcttgc |
| Pri-mir-9-7 r | tctactttcggttctctagc |

**Table S2.** Primers for site-directed mutagenesis

| Number | Gene mane | Primer sequence 5>3 |
|---|---|---|
| 1 | Pri-mir-9-1 | aggggttggctgttatctttgatctagcagtatcagtgttattc |
| 2 | Pri-mir-9-4 | tgggttagtttttctcttaggtaatcctgtatgagtttatgtgatatcataaa |
| 3 | Pri-mir-9-5 | aaatactcatacagcaagataaaacataacaactcgcttccaattcc |
| 4 | Pri-mir-9-7 | acgggttagtttttctctttcgttttcttgtatgagttatgaaatatcataaag |

**Table S3**. Sequences of primers used for qRT-PCR

| Number | name | Sequence 5'>3' | Amplicon size (bp) |
|---|---|---|---|
| 1 | pri-mir-9-1 f | agttcagccctggttgg | 149 |
| 2 | pri-mir-9-1 r | tgatgcctttcaggtgtg | |
| 3 | pri-mir-9-2 f | acttggaggcgtgttg | 120 |
| 4 | pri-mir-9-2 r | gtttacgcaatgctccatac | |
| 5 | pri-mir-9-3 f | agcagaaaaccaacagtgg | 150 |
| 6 | pri-mir-9-3 r | tctcttgctgtcttccaaag | |
| 7 | pri-mir-9-4 f | tttaaggcacatgggttag | 135 |
| 8 | pri-mir-9-4 r | tcctgcctcgttccaattc | |
| 9 | pri-mir-9-5 f | cacaagacacggaggtatcc | 166 |
| 10 | pri-mir-9-5 r | tcttcgtgtcaataggcagcc | |
| 11 | pri-mir-9-6 f | acaggaggtagttgctatc | 104 |
| 12 | pri-mir-9-6 r | tgtatgcagttaaggaggac | |
| 13 | pri-mir-9-7 f | atctgaccgcacatgtgac | 146 |
| 14 | pri-mir-9-7 r | tctactttcggttctctagc | |
| 15 | *β-actin* f | cgtgctgtcttcccatcca | 176 |
| 16 | *β-actin* r | tcaccaacgtagctgtctttctg | |
| 17 | *elavl4* f | ccgtcaacaatgtcaaggtg | 147 |
| 18 | *elavl4* r | acacttgcaagacacggtca | |
| 19 | *barhl2* f | ttcggtcacgatggagcatc | 120 |
| 20 | *barhl2* r | agcagatgaggctgaggtga | |
| 21 | *dmbx1a* f | gcgaccccatcatctataccat | 105 |
| 22 | *dmbx1a* r | cgcgtggtgagatagagactga | |
| 23 | *pax2a* f | gtgacaggtcgagagatggc | 198 |
| 24 | *pax2a* r | acttaataacgcggggttgct | |
| 25 | *otpb* f | tgttggagcgatttttcagg | 107 |
| 26 | *otpb* r | ttcctgttgatcccgtttgaa | |
| 27 | *tal1/scl* f | gcacgacggcggagaca | 226 |
| 28 | *tal1/scl* r | ggaggcagtggcaagggaa | |
| 29 | *isl1* f | gcagcagcaacccaacgacaa | 224 |
| 30 | *isl1* r | ttgagcctggaccaccttcagaa | |

**Table S4**. microRNA host genes

| MicroRNA | Ensembl gene code | Host name |
|---|---|---|
| **dre-mir-9-1** | ENSDARG00000095906 | CR848047.1 |
| **dre-mir-9-2** | ENSDARG00000098512 | si:ch73-215f7.1 |
| **dre-mir-9-3** | ENSDARG00000098395 | CU929451.2 |
| **dre-mir-9-4** | ENSDARG00000095557 | BX664610.1 |
| **dre-mir-9-5** | ENSDARG00000095715 | BX465834.1 |
| **dre-mir-9-6** | ENSDARG00000101584 | CU467822.2 |
| **dre-mir-9-7** | NA | NA |

**Table S5.** smiFISH Probe sequences for pri-mir-9-1

| Probe sequence name | Probe sequence (5' to 3') |
|---|---|
| pri-mir-9-1 smiFISH_1 | CCTCCTAAGTTTCGAGCTGGACTCAGTGcactcactttctttctcaca |
| pri-mir-9-1 smiFISH_2 | CCTCCTAAGTTTCGAGCTGGACTCAGTGgaggtctttctctctctcag |
| pri-mir-9-1 smiFISH_3 | CCTCCTAAGTTTCGAGCTGGACTCAGTGcacagcaaagtgctagaatc |
| pri-mir-9-1 smiFISH_4 | CCTCCTAAGTTTCGAGCTGGACTCAGTGgatttaccctcaagttgttg |
| pri-mir-9-1 smiFISH_5 | CCTCCTAAGTTTCGAGCTGGACTCAGTGctagttgctttagttttccg |
| pri-mir-9-1 smiFISH_6 | CCTCCTAAGTTTCGAGCTGGACTCAGTGaatagggggaaaagaccct |
| pri-mir-9-1 smiFISH_7 | CCTCCTAAGTTTCGAGCTGGACTCAGTGcgcaaaggctgggatttatt |
| pri-mir-9-1 smiFISH_8 | CCTCCTAAGTTTCGAGCTGGACTCAGTGgagagatggcttcaaacgca |
| pri-mir-9-1 smiFISH_9 | CCTCCTAAGTTTCGAGCTGGACTCAGTGattgcaggctgatcattctc |
| pri-mir-9-1 smiFISH_10 | CCTCCTAAGTTTCGAGCTGGACTCAGTGttcttgttgtctctgtctta |
| pri-mir-9-1 smiFISH_11 | CCTCCTAAGTTTCGAGCTGGACTCAGTGagatgcaaagagtccattca |
| pri-mir-9-1 smiFISH_12 | CCTCCTAAGTTTCGAGCTGGACTCAGTGaaccagcctgaactggttta |
| pri-mir-9-1 smiFISH_13 | CCTCCTAAGTTTCGAGCTGGACTCAGTGctatgacttcaccagtctgt |
| pri-mir-9-1 smiFISH_14 | CCTCCTAAGTTTCGAGCTGGACTCAGTGctcttgattgggtagcttaa |
| pri-mir-9-1 smiFISH_15 | CCTCCTAAGTTTCGAGCTGGACTCAGTGtgatcacagtgctgagactg |
| pri-mir-9-1 smiFISH_16 | CCTCCTAAGTTTCGAGCTGGACTCAGTGactgacgtaggttatctgta |
| pri-mir-9-1 smiFISH_17 | CCTCCTAAGTTTCGAGCTGGACTCAGTGggctgaactgcaacctgaaa |
| pri-mir-9-1 smiFISH_18 | CCTCCTAAGTTTCGAGCTGGACTCAGTGctttcaggtgtgtaatggga |
| pri-mir-9-1 smiFISH_19 | CCTCCTAAGTTTCGAGCTGGACTCAGTGagtacagatctcaaggtggg |
| pri-mir-9-1 smiFISH_20 | CCTCCTAAGTTTCGAGCTGGACTCAGTGagaggagaagaggagagggg |
| pri-mir-9-1 smiFISH_21 | CCTCCTAAGTTTCGAGCTGGACTCAGTGgagagagggagacttgggag |
| pri-mir-9-1 smiFISH_22 | CCTCCTAAGTTTCGAGCTGGACTCAGTGtggcttgctctcacaataaa |
| pri-mir-9-1 smiFISH_23 | CCTCCTAAGTTTCGAGCTGGACTCAGTGagcactgatggaggccattt |
| pri-mir-9-1 smiFISH_24 | CCTCCTAAGTTTCGAGCTGGACTCAGTGaaaatgctgcttaccttcca |
| pri-mir-9-1 smiFISH_25 | CCTCCTAAGTTTCGAGCTGGACTCAGTGggaggagataatcacagctt |
| pri-mir-9-1 smiFISH_26 | CCTCCTAAGTTTCGAGCTGGACTCAGTGagcaatgagagagactgggt |
| pri-mir-9-1 smiFISH_27 | CCTCCTAAGTTTCGAGCTGGACTCAGTGacgcaccttttacaatagca |
| pri-mir-9-1 smiFISH_28 | CCTCCTAAGTTTCGAGCTGGACTCAGTGtagatttgtgaggaggggtg |
| pri-mir-9-1 smiFISH_29 | CCTCCTAAGTTTCGAGCTGGACTCAGTGagtgctgactactctaatgt |
| pri-mir-9-1 smiFISH_30 | CCTCCTAAGTTTCGAGCTGGACTCAGTGattccgcgtattaaacgctt |
| pri-mir-9-1 smiFISH_31 | CCTCCTAAGTTTCGAGCTGGACTCAGTGccctacggagagtgaaatga |
| pri-mir-9-1 smiFISH_32 | CCTCCTAAGTTTCGAGCTGGACTCAGTGcattcagcttctccgtatcg |
| pri-mir-9-1 smiFISH_33 | CCTCCTAAGTTTCGAGCTGGACTCAGTGcactccagaatgttgtcttg |
| pri-mir-9-1 smiFISH_34 | CCTCCTAAGTTTCGAGCTGGACTCAGTGaaatatgcgcgtgcgagctg |
| pri-mir-9-1 smiFISH_35 | CCTCCTAAGTTTCGAGCTGGACTCAGTGattcaggaccgatcgtaatg |
| pri-mir-9-1 smiFISH_36 | CCTCCTAAGTTTCGAGCTGGACTCAGTGcctatcctcgagcatcgatg |

**Table S6.** smiFISH Probe sequences for pri-mir-9-4

| Probe sequence name | Probe sequence (5' to 3') |
|---|---|
| pri-mir-9-4 smiFISH_1 | CCTCCTAAGTTTCGAGCTGGACTCAGTGctatcgatacccttgtggaa |
| pri-mir-9-4 smiFISH_2 | CCTCCTAAGTTTCGAGCTGGACTCAGTGtggaaacgttgcgtttggtt |
| pri-mir-9-4 smiFISH_3 | CCTCCTAAGTTTCGAGCTGGACTCAGTGacactgtcaagacactgcta |
| pri-mir-9-4 smiFISH_4 | CCTCCTAAGTTTCGAGCTGGACTCAGTGatggttagcatttgttcgtg |
| pri-mir-9-4 smiFISH_5 | CCTCCTAAGTTTCGAGCTGGACTCAGTGatgcacctcaaaaaactgct |
| pri-mir-9-4 smiFISH_6 | CCTCCTAAGTTTCGAGCTGGACTCAGTGcagtatgtaggcgcttaagg |
| pri-mir-9-4 smiFISH_7 | CCTCCTAAGTTTCGAGCTGGACTCAGTGtcttttatgtttggagctgg |
| pri-mir-9-4 smiFISH_8 | CCTCCTAAGTTTCGAGCTGGACTCAGTGgcatcgcagttgtttacaat |
| pri-mir-9-4 smiFISH_9 | CCTCCTAAGTTTCGAGCTGGACTCAGTGctcttgcgtttgcactttta |
| pri-mir-9-4 smiFISH_10 | CCTCCTAAGTTTCGAGCTGGACTCAGTGtagtctaggtgttattgcgc |
| pri-mir-9-4 smiFISH_11 | CCTCCTAAGTTTCGAGCTGGACTCAGTGtacgctcatattatgtgcgt |
| pri-mir-9-4 smiFISH_12 | CCTCCTAAGTTTCGAGCTGGACTCAGTGcacacacacatacacgtggg |
| pri-mir-9-4 smiFISH_13 | CCTCCTAAGTTTCGAGCTGGACTCAGTGatacagtctcactcactctt |
| pri-mir-9-4 smiFISH_14 | CCTCCTAAGTTTCGAGCTGGACTCAGTGatggagctattgcattcact |
| pri-mir-9-4 smiFISH_15 | CCTCCTAAGTTTCGAGCTGGACTCAGTGcagatgcagcttctgttgtg |
| pri-mir-9-4 smiFISH_16 | CCTCCTAAGTTTCGAGCTGGACTCAGTGaaaactaacccatgtgcctt |
| pri-mir-9-4 smiFISH_17 | CCTCCTAAGTTTCGAGCTGGACTCAGTGtaatctcaaggtctacgggg |
| pri-mir-9-4 smiFISH_18 | CCTCCTAAGTTTCGAGCTGGACTCAGTGtcaaacgacagcactcacaa |
| pri-mir-9-4 smiFISH_19 | CCTCCTAAGTTTCGAGCTGGACTCAGTGacatccagaacatccatttc |
| pri-mir-9-4 smiFISH_20 | CCTCCTAAGTTTCGAGCTGGACTCAGTGccatccagcgttttacaaaa |
| pri-mir-9-4 smiFISH_21 | CCTCCTAAGTTTCGAGCTGGACTCAGTGaattcaaagcgctccgaagt |
| pri-mir-9-4 smiFISH_22 | CCTCCTAAGTTTCGAGCTGGACTCAGTGggattcgatgctctttgtag |
| pri-mir-9-4 smiFISH_23 | CCTCCTAAGTTTCGAGCTGGACTCAGTGggaaacgcattaacgcagcg |
| pri-mir-9-4 smiFISH_24 | CCTCCTAAGTTTCGAGCTGGACTCAGTGacagcttttggtattccagt |
| pri-mir-9-4 smiFISH_25 | CCTCCTAAGTTTCGAGCTGGACTCAGTGacgtaaaatccacgtcttcc |
| pri-mir-9-4 smiFISH_26 | CCTCCTAAGTTTCGAGCTGGACTCAGTGtatgtagccggaggcagatt |
| pri-mir-9-4 smiFISH_27 | CCTCCTAAGTTTCGAGCTGGACTCAGTGcgtactgccagcgtactaac |
| pri-mir-9-4 smiFISH_28 | CCTCCTAAGTTTCGAGCTGGACTCAGTGgtatccgcctgaataccaag |
| pri-mir-9-4 smiFISH_29 | CCTCCTAAGTTTCGAGCTGGACTCAGTGcttgagaatgatcgcgaagt |
| pri-mir-9-4 smiFISH_30 | CCTCCTAAGTTTCGAGCTGGACTCAGTGggaatgttgcccagcacaag |
| pri-mir-9-4 smiFISH_31 | CCTCCTAAGTTTCGAGCTGGACTCAGTGacatcatcatcacgtctatg |
| pri-mir-9-4 smiFISH_32 | CCTCCTAAGTTTCGAGCTGGACTCAGTGgtcatcccgaattcatacat |
| pri-mir-9-4 smiFISH_33 | CCTCCTAAGTTTCGAGCTGGACTCAGTGaacacctcgatgaaccgatt |
| pri-mir-9-4 smiFISH_34 | CCTCCTAAGTTTCGAGCTGGACTCAGTGagagatccgaccaatcggtt |
| pri-mir-9-4 smiFISH_35 | CCTCCTAAGTTTCGAGCTGGACTCAGTGtatatgagaaccacgtgacc |

**Table S7.** smiFISH Probe sequences for pri-mir-9-5

| Probe sequence name | Probe sequence (5' to 3') |
|---|---|
| pri-mir-9-5 smiFISH_1 | CCTCCTAAGTTTCGAGCTGGACTCAGTGacggaggagatcggtaatgc |
| pri-mir-9-5 smiFISH_2 | CCTCCTAAGTTTCGAGCTGGACTCAGTGttagttttggccgagtggtg |
| pri-mir-9-5 smiFISH_3 | CCTCCTAAGTTTCGAGCTGGACTCAGTGactgaatggagggatcgctc |
| pri-mir-9-5 smiFISH_4 | CCTCCTAAGTTTCGAGCTGGACTCAGTGataacatcgcgtttgtttcg |
| pri-mir-9-5 smiFISH_5 | CCTCCTAAGTTTCGAGCTGGACTCAGTGaggaaaaaaagtgcacgcgg |
| pri-mir-9-5 smiFISH_6 | CCTCCTAAGTTTCGAGCTGGACTCAGTGgaggggaaaaatccgattcc |
| pri-mir-9-5 smiFISH_7 | CCTCCTAAGTTTCGAGCTGGACTCAGTGgaaccagcagcgagattata |
| pri-mir-9-5 smiFISH_8 | CCTCCTAAGTTTCGAGCTGGACTCAGTGtggcagcagcgatagatatc |
| pri-mir-9-5 smiFISH_9 | CCTCCTAAGTTTCGAGCTGGACTCAGTGgctcctttggctctattaaa |
| pri-mir-9-5 smiFISH_10 | CCTCCTAAGTTTCGAGCTGGACTCAGTGttacctcccaatctcacaaa |
| pri-mir-9-5 smiFISH_11 | CCTCCTAAGTTTCGAGCTGGACTCAGTGtacaggccatcatttgcatt |
| pri-mir-9-5 smiFISH_12 | CCTCCTAAGTTTCGAGCTGGACTCAGTGcctgtcgaataagaagcagt |
| pri-mir-9-5 smiFISH_13 | CCTCCTAAGTTTCGAGCTGGACTCAGTGtatgagagttggagtgaggg |
| pri-mir-9-5 smiFISH_14 | CCTCCTAAGTTTCGAGCTGGACTCAGTGaccggagcagttaacgttaa |
| pri-mir-9-5 smiFISH_15 | CCTCCTAAGTTTCGAGCTGGACTCAGTGtggatatgctcgcaaccaaa |
| pri-mir-9-5 smiFISH_16 | CCTCCTAAGTTTCGAGCTGGACTCAGTGggcaatggacttcacaattg |
| pri-mir-9-5 smiFISH_17 | CCTCCTAAGTTTCGAGCTGGACTCAGTGtccaagtggagaagttgcta |
| pri-mir-9-5 smiFISH_18 | CCTCCTAAGTTTCGAGCTGGACTCAGTGctccgtgtcttgtgtaagaa |
| pri-mir-9-5 smiFISH_19 | CCTCCTAAGTTTCGAGCTGGACTCAGTGcacaacagtaatcccaggat |
| pri-mir-9-5 smiFISH_20 | CCTCCTAAGTTTCGAGCTGGACTCAGTGgataacaactcgcttccaat |
| pri-mir-9-5 smiFISH_21 | CCTCCTAAGTTTCGAGCTGGACTCAGTGctcatacagctagataacca |
| pri-mir-9-5 smiFISH_22 | CCTCCTAAGTTTCGAGCTGGACTCAGTGgaggcagtttttactttcgg |
| pri-mir-9-5 smiFISH_23 | CCTCCTAAGTTTCGAGCTGGACTCAGTGcagccagagtttcgatgagc |
| pri-mir-9-5 smiFISH_24 | CCTCCTAAGTTTCGAGCTGGACTCAGTGagctactgacacaggctata |
| pri-mir-9-5 smiFISH_25 | CCTCCTAAGTTTCGAGCTGGACTCAGTGtctaacatgttgtgtagcca |
| pri-mir-9-5 smiFISH_26 | CCTCCTAAGTTTCGAGCTGGACTCAGTGttatagcaccaagtgaagcc |
| pri-mir-9-5 smiFISH_27 | CCTCCTAAGTTTCGAGCTGGACTCAGTGcccgcgagctatgagaaata |
| pri-mir-9-5 smiFISH_28 | CCTCCTAAGTTTCGAGCTGGACTCAGTGtttgtttgcttgagttaccg |
| pri-mir-9-5 smiFISH_29 | CCTCCTAAGTTTCGAGCTGGACTCAGTGttcctccacaaggaaaagga |
| pri-mir-9-5 smiFISH_30 | CCTCCTAAGTTTCGAGCTGGACTCAGTGccagtctattcgtctatttg |
| pri-mir-9-5 smiFISH_31 | CCTCCTAAGTTTCGAGCTGGACTCAGTGttcacgtgtcagtttcagta |
| pri-mir-9-5 smiFISH_32 | CCTCCTAAGTTTCGAGCTGGACTCAGTGaatagtgagcacgtgggttc |
| pri-mir-9-5 smiFISH_33 | CCTCCTAAGTTTCGAGCTGGACTCAGTGcaaaggcgaccgagctgaaa |
| pri-mir-9-5 smiFISH_34 | CCTCCTAAGTTTCGAGCTGGACTCAGTGatttctgtttgggatgccag |
| pri-mir-9-5 smiFISH_35 | CCTCCTAAGTTTCGAGCTGGACTCAGTGgctctctctatatgtcctaa |

**Table S8.** smiFISH Probe sequences for *her6*

| **Probe sequence name** | Probe sequence (5' to 3') |
|---|---|
| her6 smiFISH_1 | CCTCCTAAGTTTCGAGCTGGACTCAGTGtccatgatatcggcaggcat |
| her6 smiFISH_2 | CCTCCTAAGTTTCGAGCTGGACTCAGTGgaccggagaagaggagtttt |
| her6 smiFISH_3 | CCTCCTAAGTTTCGAGCTGGACTCAGTGtgggtttatcaggtgtagtg |
| her6 smiFISH_4 | CCTCCTAAGTTTCGAGCTGGACTCAGTGagactttctgtgttccgaag |
| her6 smiFISH_5 | CCTCCTAAGTTTCGAGCTGGACTCAGTGtcttttctccataatgggtt |
| her6 smiFISH_6 | CCTCCTAAGTTTCGAGCTGGACTCAGTGtttcgttgattctcgctctt |
| her6 smiFISH_7 | CCTCCTAAGTTTCGAGCTGGACTCAGTGattaacgttttcagctgacc |
| her6 smiFISH_8 | CCTCCTAAGTTTCGAGCTGGACTCAGTGatcttttttcagagcatcca |
| her6 smiFISH_9 | CCTCCTAAGTTTCGAGCTGGACTCAGTGggctttctcaagtttagagt |
| her6 smiFISH_10 | CCTCCTAAGTTTCGAGCTGGACTCAGTGtcactgtcatctccaggatg |
| her6 smiFISH_11 | CCTCCTAAGTTTCGAGCTGGACTCAGTGcgctgcatgtttctgagatg |
| her6 smiFISH_12 | CCTCCTAAGTTTCGAGCTGGACTCAGTGtttagggcagcggtcatttg |
| her6 smiFISH_13 | CCTCCTAAGTTTCGAGCTGGACTCAGTGttcccaagaacggtgggatc |
| her6 smiFISH_14 | CCTCCTAAGTTTCGAGCTGGACTCAGTGattcactgaatccagctcgg |
| her6 smiFISH_15 | CCTCCTAAGTTTCGAGCTGGACTCAGTGaaccgggtaacctcgttcat |
| her6 smiFISH_16 | CCTCCTAAGTTTCGAGCTGGACTCAGTGtgttaaccccttcacatgtg |
| her6 smiFISH_17 | CCTCCTAAGTTTCGAGCTGGACTCAGTGcgttgatctgtgtcatgcag |
| her6 smiFISH_18 | CCTCCTAAGTTTCGAGCTGGACTCAGTGtgctgtgttggatagttcat |
| her6 smiFISH_19 | CCTCCTAAGTTTCGAGCTGGACTCAGTGtgaaggatggatgaggaggc |
| her6 smiFISH_20 | CCTCCTAAGTTTCGAGCTGGACTCAGTGgggatctgaaccatgggttg |
| her6 smiFISH_21 | CCTCCTAAGTTTCGAGCTGGACTCAGTGcgctaagaggcacaacgttg |
| her6 smiFISH_22 | CCTCCTAAGTTTCGAGCTGGACTCAGTGgtcaaattggaggatgagcc |
| her6 smiFISH_23 | CCTCCTAAGTTTCGAGCTGGACTCAGTGccatatactttagttgcgtc |
| her6 smiFISH_24 | CCTCCTAAGTTTCGAGCTGGACTCAGTGttgccggcacaagctggaaa |
| her6 smiFISH_25 | CCTCCTAAGTTTCGAGCTGGACTCAGTGcaaaaaggcgaactgtccgt |
| her6 smiFISH_26 | CCTCCTAAGTTTCGAGCTGGACTCAGTGttggagcaaaggcagcgttg |
| her6 smiFISH_27 | CCTCCTAAGTTTCGAGCTGGACTCAGTGtagactggaataacagggcc |
| her6 smiFISH_28 | CCTCCTAAGTTTCGAGCTGGACTCAGTGgaaccggtgtgttggaattg |
| her6 smiFISH_29 | CCTCCTAAGTTTCGAGCTGGACTCAGTGaaacggagtctgacgtgacg |

**Table S9.** smiFISH Probe sequences for *her9*

| Probe sequence name | Probe sequence (5' to 3') |
|---|---|
| *her9* smiFISH _1 | CCTCCTAAGTTTCGAGCTGGACTCAGTGcttctccatattatcggctg |
| *her9* smiFISH _2 | CCTCCTAAGTTTCGAGCTGGACTCAGTGcagcaataggtgatgctgtc |
| *her9* smiFISH _3 | CCTCCTAAGTTTCGAGCTGGACTCAGTGgcttgtcaggagtatgagat |
| *her9* smiFISH _4 | CCTCCTAAGTTTCGAGCTGGACTCAGTGagactttctatgctcgctgg |
| *her9* smiFISH _5 | CCTCCTAAGTTTCGAGCTGGACTCAGTGgcttttccatgattggcttt |
| *her9* smiFISH _6 | CCTCCTAAGTTTCGAGCTGGACTCAGTGaaggctctcgttgattctcg |
| *her9* smiFISH _7 | CCTCCTAAGTTTCGAGCTGGACTCAGTGgaatgagagtcttcagctgc |
| *her9* smiFISH _8 | CCTCCTAAGTTTCGAGCTGGACTCAGTGgctatctttttaagagcat |
| *her9* smiFISH _9 | CCTCCTAAGTTTCGAGCTGGACTCAGTGtctccaatttagagtgtctg |
| *her9* smiFISH _10 | CCTCCTAAGTTTCGAGCTGGACTCAGTGgtcatctccagaatatcagc |
| *her9* smiFISH _11 | CCTCCTAAGTTTCGAGCTGGACTCAGTGtaaattgcgcaggtgcttga |
| *her9* smiFISH _12 | CCTCCTAAGTTTCGAGCTGGACTCAGTGaaggctgcgctcatctgaac |
| *her9* smiFISH _13 | CCTCCTAAGTTTCGAGCTGGACTCAGTGtacttgctgaggacgtttgt |
| *her9* smiFISH _14 | CCTCCTAAGTTTCGAGCTGGACTCAGTGcatgcactcgttgaatcctg |
| *her9* smiFISH _15 | CCTCCTAAGTTTCGAGCTGGACTCAGTGagagaaatcgagtcacctcg |
| *her9* smiFISH _16 | CCTCCTAAGTTTCGAGCTGGACTCAGTGtctgacctctgtattcactc |
| *her9* smiFISH _17 | CCTCCTAAGTTTCGAGCTGGACTCAGTGacaggtggttaagaagtcgc |
| *her9* smiFISH _18 | CCTCCTAAGTTTCGAGCTGGACTCAGTGcatcatctgtcccatacaac |
| *her9* smiFISH _19 | CCTCCTAAGTTTCGAGCTGGACTCAGTGcaggctgagggtagttcatg |
| *her9* smiFISH _20 | CCTCCTAAGTTTCGAGCTGGACTCAGTGagccaaatgagcctgttgag |
| *her9* smiFISH _21 | CCTCCTAAGTTTCGAGCTGGACTCAGTGgaagctgcacgtgaagaggc |
| *her9* smiFISH _22 | CCTCCTAAGTTTCGAGCTGGACTCAGTGccgttgatgggtaacgttga |
| *her9* smiFISH _23 | CCTCCTAAGTTTCGAGCTGGACTCAGTGtgagtttggaacccattgag |
| *her9* smiFISH _24 | CCTCCTAAGTTTCGAGCTGGACTCAGTGtggtgagaccgcttctgaag |
| *her9* smiFISH _25 | CCTCCTAAGTTTCGAGCTGGACTCAGTGagctggaatcctccaaagac |
| *her9* smiFISH _26 | CCTCCTAAGTTTCGAGCTGGACTCAGTGaaaagcaaactgtccgtccg |
| *her9* smiFISH _27 | CCTCCTAAGTTTCGAGCTGGACTCAGTGcaaacgctgggttggggata |
| *her9* smiFISH _28 | CCTCCTAAGTTTCGAGCTGGACTCAGTGaatgaccggagttgtggcag |
| *her9* smiFISH _29 | CCTCCTAAGTTTCGAGCTGGACTCAGTGcgcttgcgtttgcgtacaag |
| *her9* smiFISH _30 | CCTCCTAAGTTTCGAGCTGGACTCAGTGactggcgttgacagtcactg |
| *her9* smiFISH _31 | CCTCCTAAGTTTCGAGCTGGACTCAGTGcattccctggacaggagatg |
| *her9* smiFISH _32 | CCTCCTAAGTTTCGAGCTGGACTCAGTGcttgtggaacgcccgagaag |
| *her9* smiFISH _33 | CCTCCTAAGTTTCGAGCTGGACTCAGTGctgacaccaacgggactgac |

**Table S10.** smiFISH Probe sequences for *neurog1*

| Probe sequence name | Probe sequence (5' to 3') |
|---|---|
| *neurog1* smiFISH_1 | CCTCCTAAGTTTCGAGCTGGACTCAGTGtgctcttaaccctcaatcag |
| *neurog1* smiFISH_2 | CCTCCTAAGTTTCGAGCTGGACTCAGTGaagataatggcacgcgtctg |
| *neurog1* smiFISH_3 | CCTCCTAAGTTTCGAGCTGGACTCAGTGatcctgcagatagtttgtgt |
| *neurog1* smiFISH_4 | CCTCCTAAGTTTCGAGCTGGACTCAGTGgagatgcttgaggttttgca |
| *neurog1* smiFISH_5 | CCTCCTAAGTTTCGAGCTGGACTCAGTGtgataaccttattggtgggc |
| *neurog1* smiFISH_6 | CCTCCTAAGTTTCGAGCTGGACTCAGTGcggagtatacgatctccatt |
| *neurog1* smiFISH_7 | CCTCCTAAGTTTCGAGCTGGACTCAGTGtagtcacagcttgaggtttc |
| *neurog1* smiFISH_8 | CCTCCTAAGTTTCGAGCTGGACTCAGTGatcatccgtgtgcgaaaagg |
| *neurog1* smiFISH_9 | CCTCCTAAGTTTCGAGCTGGACTCAGTGtggagacgcaggtggttttc |
| *neurog1* smiFISH_10 | CCTCCTAAGTTTCGAGCTGGACTCAGTGttcttcttcacgacgtgcac |
| *neurog1* smiFISH_11 | CCTCCTAAGTTTCGAGCTGGACTCAGTGttaaggttgtgcatcctgtt |
| *neurog1* smiFISH_12 | CCTCCTAAGTTTCGAGCTGGACTCAGTGgcttctcaaagcatccaatg |
| *neurog1* smiFISH_13 | CCTCCTAAGTTTCGAGCTGGACTCAGTGtgtgtcgtcaggaaacgcag |
| *neurog1* smiFISH_14 | CCTCCTAAGTTTCGAGCTGGACTCAGTGgagtctcaattttggtcagc |
| *neurog1* smiFISH_15 | CCTCCTAAGTTTCGAGCTGGACTCAGTGatgtagttgtgagcgaagcg |
| *neurog1* smiFISH_16 | CCTCCTAAGTTTCGAGCTGGACTCAGTGatccggatggtctccgaaag |
| *neurog1* smiFISH_17 | CCTCCTAAGTTTCGAGCTGGACTCAGTGatgaagacgacgaggatgcc |
| *neurog1* smiFISH_18 | CCTCCTAAGTTTCGAGCTGGACTCAGTGggtctgagttgcagtaagac |
| *neurog1* smiFISH_19 | CCTCCTAAGTTTCGAGCTGGACTCAGTGtatccaaaatcgtccatggc |
| *neurog1* smiFISH_20 | CCTCCTAAGTTTCGAGCTGGACTCAGTGctgtacactacgtcggtttg |
| *neurog1* smiFISH_21 | CCTCCTAAGTTTCGAGCTGGACTCAGTGagatgctaggcacgaagttg |
| *neurog1* smiFISH_22 | CCTCCTAAGTTTCGAGCTGGACTCAGTGaccgacatgagaacgcttaa |
| *neurog1* smiFISH_23 | CCTCCTAAGTTTCGAGCTGGACTCAGTGtacatacttctggagattct |
| *neurog1* smiFISH_24 | CCTCCTAAGTTTCGAGCTGGACTCAGTGagtaacagtggtcttacact |
| *neurog1* smiFISH_25 | CCTCCTAAGTTTCGAGCTGGACTCAGTGgttgtgctcttggttctaat |
| *neurog1* smiFISH_26 | CCTCCTAAGTTTCGAGCTGGACTCAGTGaggcaaacagtgaataccta |
| *neurog1* smiFISH_27 | CCTCCTAAGTTTCGAGCTGGACTCAGTGtctcatttgcagactgtcat |
| *neurog1* smiFISH_28 | CCTCCTAAGTTTCGAGCTGGACTCAGTGcacgctccaaggaatgcaaa |
| *neurog1* smiFISH_29 | CCTCCTAAGTTTCGAGCTGGACTCAGTGattttctctaacggggttct |
| *neurog1* smiFISH_30 | CCTCCTAAGTTTCGAGCTGGACTCAGTGcaatctgccttgctttttaa |
| *neurog1* smiFISH_31 | CCTCCTAAGTTTCGAGCTGGACTCAGTGtgacttttcaccttggacag |
| *neurog1* smiFISH_32 | CCTCCTAAGTTTCGAGCTGGACTCAGTGtcagctttatcgctctacaa |
| *neurog1* smiFISH_33 | CCTCCTAAGTTTCGAGCTGGACTCAGTGaactgatttttcacgctcgt |
| *neurog1* smiFISH_34 | CCTCCTAAGTTTCGAGCTGGACTCAGTGttcagtctattgtcacagcg |
| *neurog1* smiFISH_35 | CCTCCTAAGTTTCGAGCTGGACTCAGTGcataaggccagatctttgtc |
| *neurog1* smiFISH_36 | CCTCCTAAGTTTCGAGCTGGACTCAGTGtttcttcgggtcaaaataca |

| | |
|---|---|
| *neurog1* smiFISH_37 | CCTCCTAAGTTTCGAGCTGGACTCAGTGggatcagttggacagatgag |
| *neurog1* smiFISH_38 | CCTCCTAAGTTTCGAGCTGGACTCAGTGcatgagagctggttaactgt |
| *neurog1* smiFISH_39 | CCTCCTAAGTTTCGAGCTGGACTCAGTGagaaaagtggtgggaaagcc |
| *neurog1* smiFISH_40 | CCTCCTAAGTTTCGAGCTGGACTCAGTGcgtacaaacatgtttgcacc |

**Table S11.** smiFISH Probe sequences for *atoh1*

| **Probe sequence name** | **Probe sequence (5' to 3')** |
|---|---|
| *atoh1a* smiFISH_1 | CCTCCTAAGTTTCGAGCTGGACTCAGTGggtgaaggggatttctttac |
| *atoh1a* smiFISH_2 | CCTCCTAAGTTTCGAGCTGGACTCAGTGagatttctcttcttacacct |
| *atoh1a* smiFISH_3 | CCTCCTAAGTTTCGAGCTGGACTCAGTGtacagggacggatattttgc |
| *atoh1a* smiFISH_4 | CCTCCTAAGTTTCGAGCTGGACTCAGTGttgggaggaaagtttgtggc |
| *atoh1a* smiFISH_5 | CCTCCTAAGTTTCGAGCTGGACTCAGTGatccattctgttggtttgtg |
| *atoh1a* smiFISH_6 | CCTCCTAAGTTTCGAGCTGGACTCAGTGttcaaccacctctcttgtat |
| *atoh1a* smiFISH_7 | CCTCCTAAGTTTCGAGCTGGACTCAGTGaagctcgaatgctggacgtc |
| *atoh1a* smiFISH_8 | CCTCCTAAGTTTCGAGCTGGACTCAGTGtgaagttggaggcagaggac |
| *atoh1a* smiFISH_9 | CCTCCTAAGTTTCGAGCTGGACTCAGTGcgtactttgacaggactctt |
| *atoh1a* smiFISH_10 | CCTCCTAAGTTTCGAGCTGGACTCAGTGgttggtggatttgctggatg |
| *atoh1a* smiFISH_11 | CCTCCTAAGTTTCGAGCTGGACTCAGTGttcaatccgtgcattcttcg |
| *atoh1a* smiFISH_12 | CCTCCTAAGTTTCGAGCTGGACTCAGTGcaaaggctgggatgacactg |
| *atoh1a* smiFISH_13 | CCTCCTAAGTTTCGAGCTGGACTCAGTGttggagagtttcttgtcgtt |
| *atoh1a* smiFISH_14 | CCTCCTAAGTTTCGAGCTGGACTCAGTGttgatgtagatctgggccat |
| *atoh1a* smiFISH_15 | CCTCCTAAGTTTCGAGCTGGACTCAGTGctgtagtaagtcggacaggg |
| *atoh1a* smiFISH_16 | CCTCCTAAGTTTCGAGCTGGACTCAGTGtttctaacacgttggcatgc |
| *atoh1a* smiFISH_17 | CCTCCTAAGTTTCGAGCTGGACTCAGTGctcgtactggtacgggtaac |
| *atoh1a* smiFISH_18 | CCTCCTAAGTTTCGAGCTGGACTCAGTGgtcttgctccatgaaagagt |
| *atoh1a* smiFISH_19 | CCTCCTAAGTTTCGAGCTGGACTCAGTGcgaaccagacttgctcgttc |
| *atoh1a* smiFISH_20 | CCTCCTAAGTTTCGAGCTGGACTCAGTGaccgaggcgagtctttactg |
| *atoh1a* smiFISH_21 | CCTCCTAAGTTTCGAGCTGGACTCAGTGcgagtgaggcgagaactctc |
| *atoh1a* smiFISH_22 | CCTCCTAAGTTTCGAGCTGGACTCAGTGgtttcgtctgagtcactgaa |
| *atoh1a* smiFISH_23 | CCTCCTAAGTTTCGAGCTGGACTCAGTGgacagctcgtcttcactctg |
| *atoh1a* smiFISH_24 | CCTCCTAAGTTTCGAGCTGGACTCAGTGtttcttaaaaagcgcggcgc |
| *atoh1a* smiFISH_25 | CCTCCTAAGTTTCGAGCTGGACTCAGTGtgacacgattgagggacagt |
| *atoh1a* smiFISH_26 | CCTCCTAAGTTTCGAGCTGGACTCAGTGaagcaacccattacaaagcc |
| *atoh1a* smiFISH_27 | CCTCCTAAGTTTCGAGCTGGACTCAGTGtctattggcgtcagacacaa |
| *atoh1a* smiFISH_28 | CCTCCTAAGTTTCGAGCTGGACTCAGTGgtcaatgtcacgagaaggca |
| *atoh1a* smiFISH_29 | CCTCCTAAGTTTCGAGCTGGACTCAGTGttatgcatgacattcgagcc |
| *atoh1a* smiFISH_30 | CCTCCTAAGTTTCGAGCTGGACTCAGTGgtctcaaaacagtttgcgga |
| *atoh1a* smiFISH_31 | CCTCCTAAGTTTCGAGCTGGACTCAGTGaggtccatgacaatcatgtg |
| *atoh1a* smiFISH_32 | CCTCCTAAGTTTCGAGCTGGACTCAGTGgtcgcaaattgttacagggt |
| *atoh1a* smiFISH_33 | CCTCCTAAGTTTCGAGCTGGACTCAGTGccgtttttaaagtgcaagt |
| *atoh1a* smiFISH_34 | CCTCCTAAGTTTCGAGCTGGACTCAGTGgaatatttctcaagcctacg |
| *atoh1a* smiFISH_35 | CCTCCTAAGTTTCGAGCTGGACTCAGTGcacatcatttcttgttccat |
| *atoh1a* smiFISH_36 | CCTCCTAAGTTTCGAGCTGGACTCAGTGattggcattacttctacata |
| *atoh1a* smiFISH_37 | CCTCCTAAGTTTCGAGCTGGACTCAGTGgcagaacacaacttctttgc |
| *atoh1a* smiFISH_38 | CCTCCTAAGTTTCGAGCTGGACTCAGTGatagcttacttcagctacag |
| *atoh1a* smiFISH_39 | CCTCCTAAGTTTCGAGCTGGACTCAGTGgctcatttccaatgtaacac |

**Table S12.** smiFISH Probe sequences for *ascl1*

| Probe sequence name | Probe sequence (5' to 3') |
|---|---|
| *ascl1a* smiFISH_1 | CCTCCTAAGTTTCGAGCTGGACTCAGTGtctcgacctgttctgagttg |
| *ascl1a* smiFISH_2 | CCTCCTAAGTTTCGAGCTGGACTCAGTGgttcacgtgtggcttcaatg |
| *ascl1a* smiFISH_3 | CCTCCTAAGTTTCGAGCTGGACTCAGTGaaagttttctttggactgcc |
| *ascl1a* smiFISH_4 | CCTCCTAAGTTTCGAGCTGGACTCAGTGtccatttcgcggagtcaaaa |
| *ascl1a* smiFISH_5 | CCTCCTAAGTTTCGAGCTGGACTCAGTGtggtttacgcttatttccat |
| *ascl1a* smiFISH_6 | CCTCCTAAGTTTCGAGCTGGACTCAGTGaagcaagcaggtggcatgaa |
| *ascl1a* smiFISH_7 | CCTCCTAAGTTTCGAGCTGGACTCAGTGtgagttggatgctctgagag |
| *ascl1a* smiFISH_8 | CCTCCTAAGTTTCGAGCTGGACTCAGTGttgacgcggacttgttgctg |
| *ascl1a* smiFISH_9 | CCTCCTAAGTTTCGAGCTGGACTCAGTGattgagtcttcttttgcacc |
| *ascl1a* smiFISH_10 | CCTCCTAAGTTTCGAGCTGGACTCAGTGgggtaagctgtagccgaaac |
| *ascl1a* smiFISH_11 | CCTCCTAAGTTTCGAGCTGGACTCAGTGcaaagccgttgttcacaagc |
| *ascl1a* smiFISH_12 | CCTCCTAAGTTTCGAGCTGGACTCAGTGctccattgggaacgtgttcg |
| *ascl1a* smiFISH_13 | CCTCCTAAGTTTCGAGCTGGACTCAGTGctttgctcatcttcttgttg |
| *ascl1a* smiFISH_14 | CCTCCTAAGTTTCGAGCTGGACTCAGTGgtagttttgggagatggtgg |
| *ascl1a* smiFISH_15 | CCTCCTAAGTTTCGAGCTGGACTCAGTGgccatagagttcatgtcatt |
| *ascl1a* smiFISH_16 | CCTCCTAAGTTTCGAGCTGGACTCAGTGctcatccgatgagtatgagg |
| *ascl1a* smiFISH_17 | CCTCCTAAGTTTCGAGCTGGACTCAGTGttgttcttctggactcagag |
| *ascl1a* smiFISH_18 | CCTCCTAAGTTTCGAGCTGGACTCAGTGaaaccagttggtgaagtcca |
| *ascl1a* smiFISH_19 | CCTCCTAAGTTTCGAGCTGGACTCAGTGagtttccttttacgaacgct |
| *ascl1a* smiFISH_20 | CCTCCTAAGTTTCGAGCTGGACTCAGTGccaagcgagtgctgatattt |
| *ascl1a* smiFISH_21 | CCTCCTAAGTTTCGAGCTGGACTCAGTGttgtgttcttggaggacatc |
| *ascl1a* smiFISH_22 | CCTCCTAAGTTTCGAGCTGGACTCAGTGcttggctctttgacactcgg |
| *ascl1a* smiFISH_23 | CCTCCTAAGTTTCGAGCTGGACTCAGTGatagatttcttgggcgagtg |
| *ascl1a* smiFISH_24 | CCTCCTAAGTTTCGAGCTGGACTCAGTGggtgtcgtggaaagtctttt |
| *ascl1a* smiFISH_25 | CCTCCTAAGTTTCGAGCTGGACTCAGTGcaacgtttgcttgttgttgt |
| *ascl1a* smiFISH_26 | CCTCCTAAGTTTCGAGCTGGACTCAGTGagggcaaaccttctttgatt |
| *ascl1a* smiFISH_27 | CCTCCTAAGTTTCGAGCTGGACTCAGTGagagttttgagagaggggtc |
| *ascl1a* smiFISH_28 | CCTCCTAAGTTTCGAGCTGGACTCAGTGaattcgccaagttggaagca |
| *ascl1a* smiFISH_29 | CCTCCTAAGTTTCGAGCTGGACTCAGTGcggcaggctataggtcaaaa |
| *ascl1a* smiFISH_30 | CCTCCTAAGTTTCGAGCTGGACTCAGTGctgcgatgcatttgagacta |
| *ascl1a* smiFISH_31 | CCTCCTAAGTTTCGAGCTGGACTCAGTGctccacaaccgtaaagggaa |
| *ascl1a* smiFISH_32 | CCTCCTAAGTTTCGAGCTGGACTCAGTGttgagcattacactcctcta |
| *ascl1a* smiFISH_33 | CCTCCTAAGTTTCGAGCTGGACTCAGTGataagacacgttggtgctga |
| *ascl1a* smiFISH_34 | CCTCCTAAGTTTCGAGCTGGACTCAGTGtcactctgagtcacattaca |
| *ascl1a* smiFISH_35 | CCTCCTAAGTTTCGAGCTGGACTCAGTGggacaatagctgcataacct |
| *ascl1a* smiFISH_36 | CCTCCTAAGTTTCGAGCTGGACTCAGTGgagacacaaaacacctccgt |

| | |
|---|---|
| *ascl1a* smiFISH_37 | CCTCCTAAGTTTCGAGCTGGACTCAGTGgcaaagtggaacaggcagtg |
| *ascl1a* smiFISH_38 | CCTCCTAAGTTTCGAGCTGGACTCAGTGtgactgcaacacgtaaagca |
| *ascl1a* smiFISH_39 | CCTCCTAAGTTTCGAGCTGGACTCAGTGttagcaggatggcttatcac |
| *ascl1a* smiFISH_40 | CCTCCTAAGTTTCGAGCTGGACTCAGTGttggcattcattaagagcgc |

**Table S13.** CRISPR/Cas9 Target sequences for pre-mir-9-1 and primers used to generate respective sgRNA. PAM region is Highlighted in red.

| sgRNA Number | Target sequence 5'-3' with PAM | CRISPRscan Primer 5'-3' |
|---|---|---|
| 1 | GGACGGGTGGCCGGAGGGGTTGG | taatacgactcactataGGACGGGTGGCCGGAGGGGTgttttagagctagaa |
| 2 | AAGGTGGGGTGATGCCTTTCAGG | taatacgactcactataGGGGTGGGGTGATGCCTTTCgttttagagctagaa |
| 3 | AGAGAGAGGGAGACTTGGGAGGG | taatacgactcactataGGAGAGAGGGAGACTTGGGAgttttagagctagaa |

**Table S14. The parameter values used for the mathematical model.** Highlighted in grey are the additional parameters used for the extended model. $\alpha_X$, $\mu_X$, $\alpha_h$ and $\mu_h$ parameters were used in both, adaptation and extended, models.

| **Parameter** | **Value** |
|---|---|
| $h_1$ | 3.0 |
| $h_2$ | 3.0 |
| $h_3$ | 3.0 |
| $P_1$ (low, high) | 2.0, 10.0 |
| $p_2$ | 1.0 |
| $p_3$ | 1.0 |
| $\alpha_X$ | 0.1 |
| $\mu_X$ | 0.1 |
| $\alpha_h$ | 6.0 |
| $\mu_h$ | 2.0 |
| $\alpha_Y$ | 30.0 |
| $\beta_Y$ | 100.0 |
| $\mu_Y$ | 10.0 |
